# Linking interpersonal differences in gut microbiota composition and drug biotransformation activity

**DOI:** 10.64898/2026.01.21.700809

**Authors:** Eleonora Mastrorilli, Pamela Herd, Federico E. Rey, Andrew L. Goodman, Michael Zimmermann

## Abstract

Individuals vary widely in their responses to drugs, and growing evidence implicates the gut microbiome as a contributor to this variability. While prior studies show that gut bacteria can metabolize drugs, how differences in microbial community composition influence drug metabolism remains poorly understood. Here, we characterize the biotransformation of 271 drugs by 89 gut microbial communities derived from human donors and preclinical animal models. Over 90% of tested drugs were metabolized by at least one microbiome. We identified 66 drugs exhibiting highly variable metabolism across human-derived microbiomes and several drugs whose biotransformation differed markedly between human and animal microbiomes. To enable prediction of microbiota-mediated drug metabolism, we developed and compared multiple modeling approaches based on metagenomic data. These results, together with the provided data and analytical resources contribute to a better understanding of microbiome-drug interactions and support their future integration into drug discovery, personalized prescription, and therapeutic drug monitoring.

## Introduction

The response to drugs can differ significantly among patients, and frequently, the effectiveness of drugs varies across different regions and cultural backgrounds without clear explanation^1^. Consequently, patients may experience extended periods of trial and error when adjusting dosages and drug schedules, leading to less-than-ideal treatment outcomes, unwanted drug reactions, and additional complexities^2^. A key objective of personalized medicine is to anticipate individual responses to drugs, enabling customized therapeutic approaches. Although human genome sequencing and pharmacogenomics approaches have been able to address some of these issues, interpersonal differences in drug efficacy, metabolism, and toxicity often remain challenging to predict^3^.

Growing evidence implicates the human gut microbiota in the metabolism of medical drugs^4–6^, with potential consequences on pharmacodynamics^6–10^, pharmacokinetics^11^, and drug toxicity^11–13^. Significant progress has been made to assess the biotransformation capacity of gut bacteria across a broad chemical diversity of drug compounds and specific biotransformation reactions have been successfully mapped to individual bacteria and metabolic pathways. However, despite these mechanistic insights, quantification of how the vast inter-individual differences in microbiome composition translates to interpersonal difference in microbiota drug metabolism remains challenging. This creates a significant roadblock in the use of microbiome markers to personalize drug treatments. Moreover, the lack of systematic analyses of gut microbiota drug metabolism in preclinical animal models commonly employed in drug discovery and development hampers the rational integration of gut microbial metabolism in the development of new therapeutic compounds. Furthermore, methods to understand and predict microbial community metabolism of medical drugs will have impact on microbiome interaction with diverse exogenous (xenobiotic) and endogenous small molecules.

In this study, we systematically measured the biotransformation kinetics of 271 different drugs upon exposure to 60 human gut communities and 29 gut communities derived from the most common preclinical animal species. The resulting >23,500 drug-microbiota interactions identify chemical features targeted in predictable ways by human gut microbiomes, inform predictive models, and identify compounds that are differentially metabolized in humans compared to pre-clinical animal species. These metagenomic, metabolomic, and modelling results provide a resource for the 271 approved drugs studied here, as well as other small molecules with shared chemical features.

## Results

### Gut microbiota of 60 individuals collectively biotransform 242 of 271 tested medical drugs

To assess interpersonal differences in microbiota drug metabolism, we measured the biotransformation kinetics of 271 different medical drugs by gut microbial communities collected from 60 human donors who had provided fecal samples as part of the Wisconsin Longitudinal Study^14^ (Figure 1A, Table S1). Faecal samples were cultured under anaerobic conditions for 24 hours as previously reported^4^, followed by metagenomic shotgun sequencing to determine their microbiome composition (Figure S1A, Table S2). We compared the microbiome composition of these 60 gut communities to publicly available gut microbiome metagenomes of 2803 healthy Western donors^15^ at the genus level using both Bray-Curtis^16^ and Jaccard^17^ distance measures. PERMANOVA^18^ tests using either metric were not significant in both the overall test and in 100% of 1000 subset bootstrap repetitions, demonstrating that the fecal communities from these 60 donors well represent westernized gut microbiome diversity (Figure 1B). To assay drug-metabolizing activities for each of the 60 communities, we incubated them under anaerobic culture conditions with 271 chemically diverse drugs using a previously developed combinatorial pooling scheme^4^. We collected samples at nine time points (0, 1, 2, 4, 6, 8, 12, 18, and 24 h of incubation) for drug quantification by liquid chromatography-coupled mass spectrometry (LC–MS). The analysis of 14,472 LC-MS runs captured the biotransformation kinetics of 271 drugs by 60 human gut communities in quadruplicate, plus non-bacterial controls to assess spontaneous drug degradation in complex culture media at pH 4, 5, 6, and 7 (Table S3). To identify drugs metabolized by the microbiota, we first compared drug levels at the beginning and after 24 h of the biotransformation assays. After excluding 6 drugs that could not be reliably measured, we found 242 of the remaining 265 tested drugs to be metabolized by at least one of the 60 bacterial communities. These communities were able to reduce the levels of at least 35, and as many to 206, drugs by at least 25% in two consecutive timepoints (adjusted p-value <0.1), with an average of 136 drugs metabolized per community (Figure S1B, Table S4). Only five drugs were metabolized by all communities (*i.e.*, bisacodyl, cetirizine, pranoprofen, diacetamate, and omeprazole), and 23 drugs where not metabolized by any of the communities (Figure S1C, Table S5). Notably, we found that the levels of all drugs that we had previously identified to be metabolized by isolated bacterial strains^4^ also significantly decreased upon exposure to complex microbial gut communities with the exception of three drugs (*i.e.*, eszopiclone, bicalutamide, naftopidil). Intriguingly, we identified 64 drugs that were metabolized by one or more of the tested bacterial communities, but not by the 76 individual (axenic) species and strains examined previously (Table S5)^4^. Nine out of these 64 drugs were fully converted (>90% reduction of drug levels) by at least one community (Figure S1C). These findings suggest that these drugs are either metabolized by gut bacteria present in the gut community that were not included as axenic cultures in the previous study, or that their bacterial biotransformation depends on microbial community context, for example through emerging metabolic capabilities^19^. Additionally, gut microbiomes from individual donors exhibit significant differences in their gut microbial drug metabolism capacity: while any two communities share the ability to metabolize 98±29 drugs on average, almost as many (84±58) drugs are significantly metabolized by one of these communities but not the other. For example, communities 077 and 215 collectively metabolize 196 drugs by at least 25%, but 122 of these drugs are metabolized by one of these communities but not the other. We next aimed to systematically quantify these interpersonal differences in gut microbiota drug metabolism.

**Figure 1.**
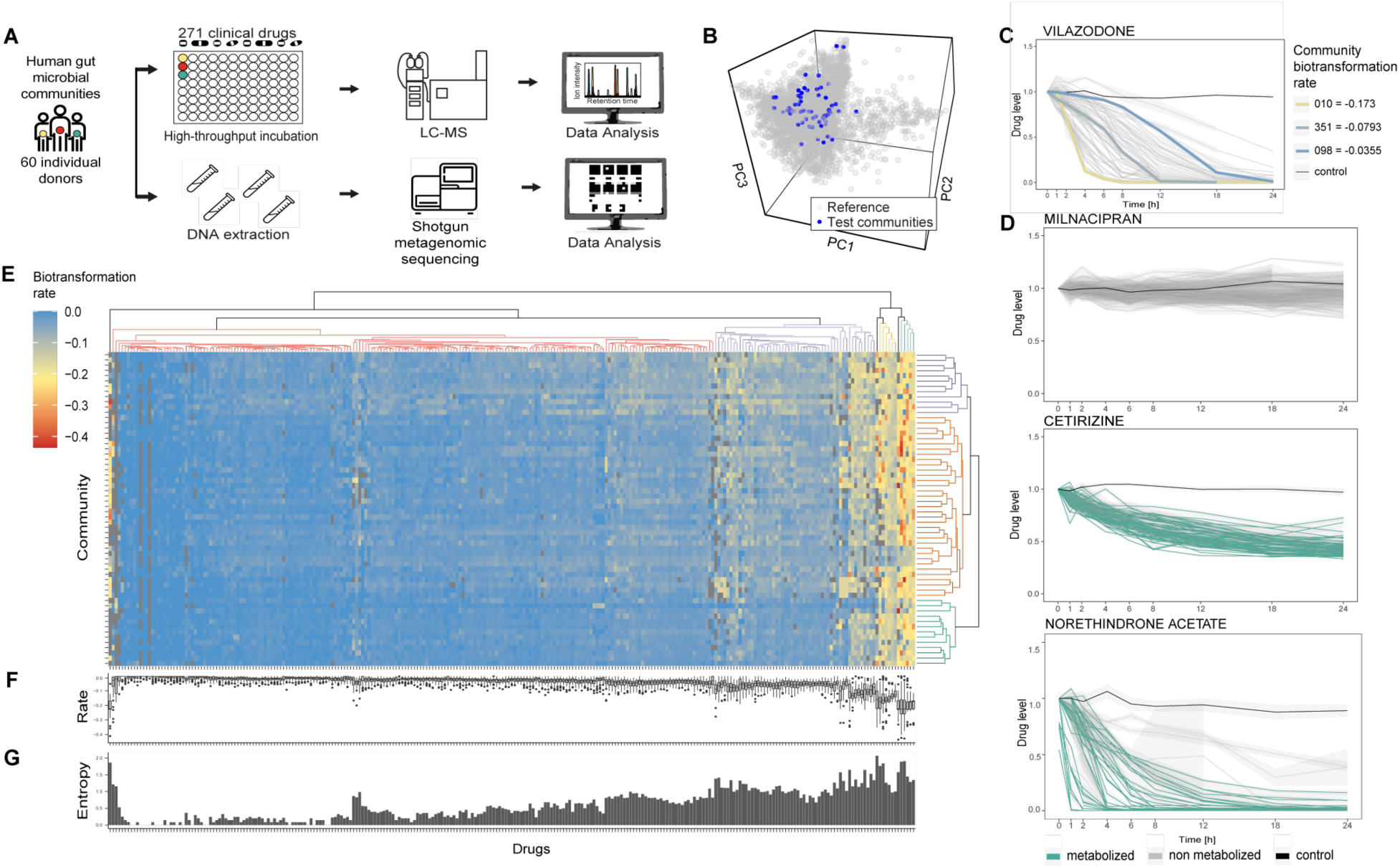
Human gut microbiota shows biotransformation activity on medical drugs. (A) Scheme of the high-throughput pipeline to assay gut microbial drug metabolism. (B) Beta diversity of the tested 60 human gut microbial communities compared to 2803 microbiomes derived from healthy Western donors (ExperimentHub^15^). (C) Biotransformation kinetics of vilazodone, which is highly community-dependent. Three example communities are highlighted in distinct colours. (D) Examples of commonly observed patterns of biotransformation: certain drugs were not metabolized by any of the communities (*e.g.,* Milnacipran, top panel); other compounds were metabolized at comparable rates across communities (*e.g.,* Cetirizine, middle panel); and other drugs show high variability in biotransformation rates across communities (*e.g.*, Norethindrone Acetate, bottom panel). (E) Heatmap of drug biotransformation rates and hierarchical clustering of communities and drugs. Grey squares indicate drug/community combinations for which it was not possible to estimate a bioransformation rate. (F) Boxplot showing the distribution of biotransformation rates for each drug across the 60 microbial communities (center line, median; box, interquartile range; whiskers, 1.5× interquartile range) (G) Shannon Entropy of the biotransformation rates for each drug across the 60 microbial communities. All drug-microbiota measurements shown (Figure 1C-G) were collected in quadruplicate.

### Gut microbial drug biotransformation depends on individuals and drugs

While the comparison of drug levels before and after microbiota incubation provides information about the biotransformation capacity of microbial communities, insights into interpersonal differences in biotransformation rates are limited. This is exemplified by vilazodone, which is fully metabolized by most of the 60 microbial communities after 24 h; however, biotransformation kinetics vary widely (Figure 1C). To systematically compare drug biotransformation rates across communities, we took advantage of the collected time-series data to model the biotransformation kinetics for each community-drug pair using a linear approximation of the initial slope of the drug concentration over time (Figure S2A-B, Table S6). The resulting biotransformation rates for the different drugs revealed several distinct patterns (Figure 1D): (i) compounds that were not metabolized by any of the communities, (ii) compounds metabolized at comparable rates across communities, and (iii) drugs showing high variability in biotransformation rates across communities. To test whether high and low drug biotransformation rates across drugs are an inherent property of a given bacterial community, we investigated whether biotransformation rates of different drugs would rank similarly across communities. This did not appear to be the case as the average correlation coefficient among two drug rankings was-0.0167± 0.151. For example, the fast-metabolizing microbial communities for sulfinyprazone are distinct from the fast-metabolizing communities for prazosin (Figure S2C).

To systematically investigate patterns in drug biotransformation rates, we hierarchically clustered them across both communities and drugs (Figure 1E), which revealed distinct clusters of bacterial communities and drugs consistent with the three patterns (i-iii) highlighted above (Figure 1D). Intriguingly, these clusters also illustrate that interpersonal differences in gut microbiota drug metabolism are drug dependent (Figure 1F), which we quantified as the Shannon Entropy^20^ of the biotransformation rates for each drug across the 60 microbial communities (Figure 1G, Table S7). Calculated entropies range between 0 and 2.0, with higher values depicting stronger interpersonal differences, quantified as higher uncertainty in biotransformation rates for a given drug between individuals’ microbiota. The five drugs showing the highest entropy were norethindrone acetate, sulfasalazine, nitrendipine, danazol, and prednisone. Altogether, these analyses suggest that biotransformation rates vary significantly between drugs and between gut microbial communities.

### Chemical features drive gut microbial drug biotransformation

We next compared the clustering of drug biotransformation rates across the 60 communities (shown in Figure 1E) to the chemical similarity between drugs, measured by Tanimoto distances of Morgan chemical fingerprints (Figure 2A, Table S8). While chemical similarities between certain drugs directly relates to their observed gut microbial biotransformation rates, the overall entanglement between biotransformation rates and chemical similarity was low (0.38) (Figure 2A), as well as the overall correlation between measured distances (PCC = 0.117, p-value = 2.03e-4, Figure S3A). This suggests that chemical similarity of entire drug molecules is likely not sufficiently granular to explain microbiota drug biotransformation. Therefore, we decided to look for chemical features of drug molecules that could be linked to biotransformation differences between microbial communities. To this end, we performed an over-representation analysis of chemical groups of the drugs according to their interpersonal variation (*i.e.*, entropy) in gut microbiota biotransformation rates (Table S9). We found that drugs showing high entropy are enriched for esters and nitro groups, which have indeed already been shown to be the target of gut bacterial biotransformation reactions^4,21,22^. Further we found enrichment of allylic oxidation sites and acetylenes, which have not yet been shown to be a target of bacterial drug biotransformation, among high-entropy drugs. On the contrary, we found that drugs with low entropies reflecting shared lack of metabolism across communities are enriched for amines including quaternary nitrogen, suggesting a possible role of positive charges in prohibiting bacterial drug metabolism. Indeed, we could confirm that drugs showing entropy below 0.66 are enriched in both quaternary nitrogen as well as positive charges at pH 7 (as computed as balanced charges from the drug structure) and that these two features are highly and significantly correlated (PCC = 0.776; pval < 2.2e-16).

**Figure 2.**
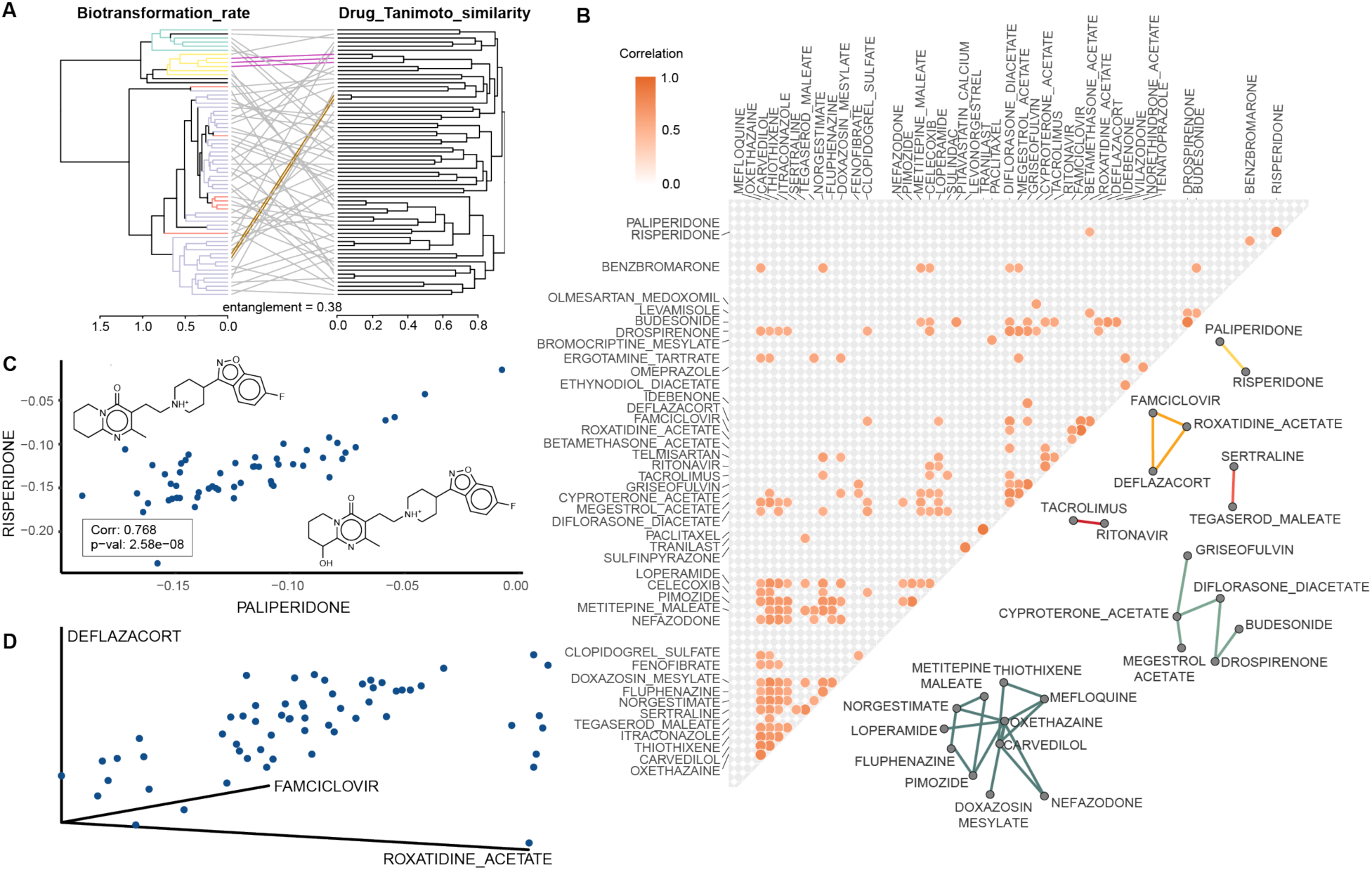
Chemical dissimilarity of the tested drugs and its relationship with observed drug biotransformation rates in gut communities. (A) Tanglegram between the hierarchical clustering of drug biotransformation rates (left) and the drug structures based on Tanimoto distances (right). For visibility purposes, only the 66 drugs (top 25%) showing the highest entropy values of biotransformation rates are represented. Overall entanglement is low and only a few examples show similar clustering between the two analyses (highlighted in golden and fuchsia). Drug slopes are coloured according to Figure 1E. (B) Pairwise correlations (PCC≥ 0.7) for the biotransformation rates of the 66 drugs (top 25%) showing the highest entropy values. 26 drugs were found to be highly correlated and could be clustered in 6 distinct clusters. (C) Scatterplot of the biotransformation rates of risperidone and paliperidone, showing both high chemical similarity and high correlation in biotransformation rate. In the box, Pearson Correlation coefficient and its significance are indicated. (D) 3D scatterplot of the biotransformation rates of deflazacort, famciclovir and roxatidine acetate, showing low chemical similarity (Figure S3B-D) but high correlation in biotransformation rates across 60 human gut communities.

To further investigate which chemical features are associated with the greatest variation in drug biotransformation rates across gut bacterial communities, we selected the 66 drugs (25%) that showed the highest entropy (Figure 1G, Table S7). For these drugs, we then compared their biotransformation rates for a given drug between communities to identify drugs showing highly correlated biotransformation across all communities. We found 26 highly correlated drugs (PCC ≥ 0.7), which form a network of six distinct clusters (Figure 2B, Table S10). One of these clusters of correlating drugs (PCC = 0.768) connects risperidone and paliperidone, two structurally related atypical antipsychotics (Tanimoto distance: 0.0783) that undergo the same gut microbial biotransformation^4^ (Figure 2C). However, other clusters connect more disparate drugs: for example, famciclovir (antiviral), roxatidine acetate (H2 blocker), and deflazacort (glucocorticoid) are metabolized at similar rates by the 60 tested communities despite minimal overall similarity of their chemical structures (Figure S3B-G). Maximum Common Substructure analysis^23^ (MCS) highlighted that all three drugs share an acetyl group (Figure S3H), that was previously shown to be hydrolysed by gut bacteria^4^. This provides a mechanistic explanation for the strong correlation of the biotransformation rates of these three drugs across the 60 different gut microbial communities despite their marked differences in chemical structure and clinical indication. Together, these analyses suggest that structural comparison of compounds with correlating biotransformation rates across communities can highlight chemical features that determine interpersonal differences in microbiome-mediated drug biotransformation (Figure S4).

### Microbiome composition drives gut microbial drug biotransformation

To understand how the compositional differences between the 60 microbial communities relate to their drug biotransformation activity, we leveraged the results of the shotgun metagenomics analysis (Figure 1A-B and Figure S1A). We first noted that overall community alpha-diversity was not strongly correlated with the number of drugs metabolized per community (PCC = 0.25, p-value 0.058, Figure S5A). Next, we compared the similarity of the 60 communities based on either their microbiome composition (measured by Bray-Curtis distance) and their drug biotransformation rates (Figure 1E). Both entanglement measurement (0.36) and overall correlation between measured distances (PCC = 0.151, p-value = 2.025e-4) showed only a weak link between the microbiota composition (as measured by beta-diversity) and the drug biotransformation activity of the gut microbial communities (Figure 3A, Figure S5B). Together, these results suggest that overall microbiome diversity alone is insufficient to explain gut microbial drug metabolism.

**Figure 3.**
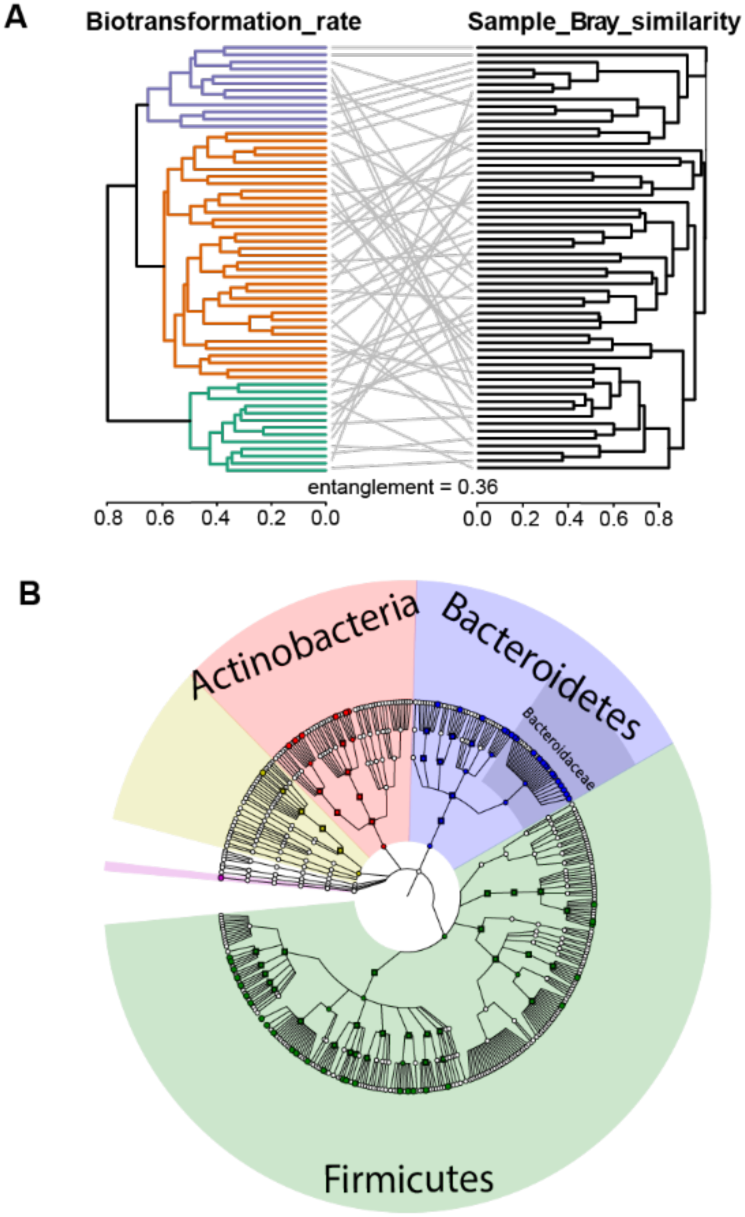
Community composition of the human gut microbiota and its relationship with observed drug biotransformation rates. (A) Tanglegram between the hierarchical clustering of communities based on their drug biotransformation rates (left) and the hierarchical clustering of communities based on their composition (Bray-Curtis similarity) (right). Overall entanglement is low and only a few examples show similar clustering for both microbiome composition and microbiota drug biotransformation. (B) Presence of 53 bacterial species that were previously assayed for the biotransformation of the 271 drugs in axenic cultures in the 60 human gut communities (Table_S16). Detected taxa are indicated as colored nodes.

Given that metabolic functions do not necessarily follow bacterial taxonomy^24^, we next used the collected metagenomic data to test the correlation between drug biotransformation rates and the abundance of 31 gut bacterial genes that have previously been experimentally validated to biotransform 21 of the tested drugs (Table S11, Table S12, Table S13, Figure S5C)^4^. We found that only the biotransformation rate of artemisinin significantly correlates with the abundance of the associated metabolic gene (SCC =-0.305, p-val = 0.0198) (Table S14, Figure S5D). Although genes encoding drug-metabolizing enzymes can serve as biomarkers to predict community activity^4,7,25,26^, these analyses suggest that for many drugs, microbiome metabolic activity will not be readily predicted by the abundance of a single gene. To identify specific microbiome features that could better explain interpersonal differences in microbiota drug metabolism, we calculated the Spearman correlation between each taxonomic feature across all communities (8 phyla, 14 classes, 19 orders, 43 families 108 genera and 370 species) and the biotransformation rates of each drug. This led to the identification of 13, 23, 63, 239, 379 and 382 significant drug biotransformation rate-taxa correlations (FDR-corrected p-value ≤ 0.05) at the phylum, class, order, family, genus, and species level, respectively (Table S15). Intriguingly, in some cases the association of an individual species with the biotransformation rates of a drug is also reflected at higher taxonomic levels (*e.g.*, norethindrone acetate significantly correlates with the abundance of *Ruminococcus torques* (SCC =-0.594, adj. p-val = 7.1e-3) and also with the family Lachnospiraceae (SCC =-0.48, adj. p-val = 0.023). However, the biotransformation rate of many drugs could only be correlated with taxonomic feature(s) at specific taxonomic levels, such as the case of mycophenolate mofetil that significantly correlated with the abundance of *Prevotella copri* (SCC =-0.486, adj.p-val = 0.026) but not with any higher taxonomic group. Noteworthy, *R. torques* and *P. copri* were previously shown to biotransform norethindrone and mycophenolate, respectively^4^, suggesting that the observed associations between specific microbiome features and community drug biotransformation result from metabolic activities of specific microbial species or taxa. Although we note that (i) correlation does not imply causation and that (ii) spurious correlations can arise in our experimental setup (especially due to the high dimensionality and compositional nature of microbiome data), this data-driven approach enriches previous knowledge of bacterial taxa involved in drug biotransformation and the observed interpersonal differences in gut microbial drug metabolism.

### Modelling gut microbial drug biotransformation activity based on microbiome composition

Because variation in microbiome biotransformation activity of certain drugs is correlated with the community abundance of specific bacterial species, we employed modelling approaches to test whether individuals’ gut microbiota drug biotransformation rates could be predicted based on microbiome composition. To this aim, we focused on the 66 drugs that displayed the strongest interpersonal differences in microbiota drug biotransformation (25% of tested drugs with highest entropy). In our first modelling approach, we generated linear models with the 53 bacterial species that were previously assayed for the biotransformation of the 66 drugs and that we detected in at least one of the 60 assayed microbial communities (Figure 3B, Table S16). Out of the 66 drugs, 50 were previously shown to be biotransformed in axenic bacterial cultures by at least one of the 53 bacterial species (Table S17). Using this set of data, we tested whether community drug biotransformation rates could be predicted by the abundance of distinct species or stepwise linear combinations thereof. As a second modelling approach, we employed generalized linear models (glm) that include the abundance of all measured taxa and of the 31 quantified genes encoding drug-metabolizing enzymes (Table S13). These models thus include bacterial species without prior experimental evidence of drug biotransformation activity (>85% of the species in the tested communities), contributions of higher taxonomic levels, potential additive multi-species effects, and relevant genes. As a third modelling approach, we developed regression models based on Random Forest (RF)^27^, which requires no formal distributional or relational assumption between the predictors and the outcome. We assessed the performance of all models by measuring Root Mean Square Error (RMSE) and Pearson Correlation Coefficient (PCC) between predicted and observed drug biotransformation rates in a nested cross-validation setting to ensure robust and unbiased performance estimates independent of the sample size^28^ (Figure 4A). In total, we built 757 single species models, 50 combined linear models, 65 glm models, and 65 RF models (after removal of entacapone, for which biotransformation rates could be estimated for less than ten microbial gut communities) (Table S18).

**Figure 4.**
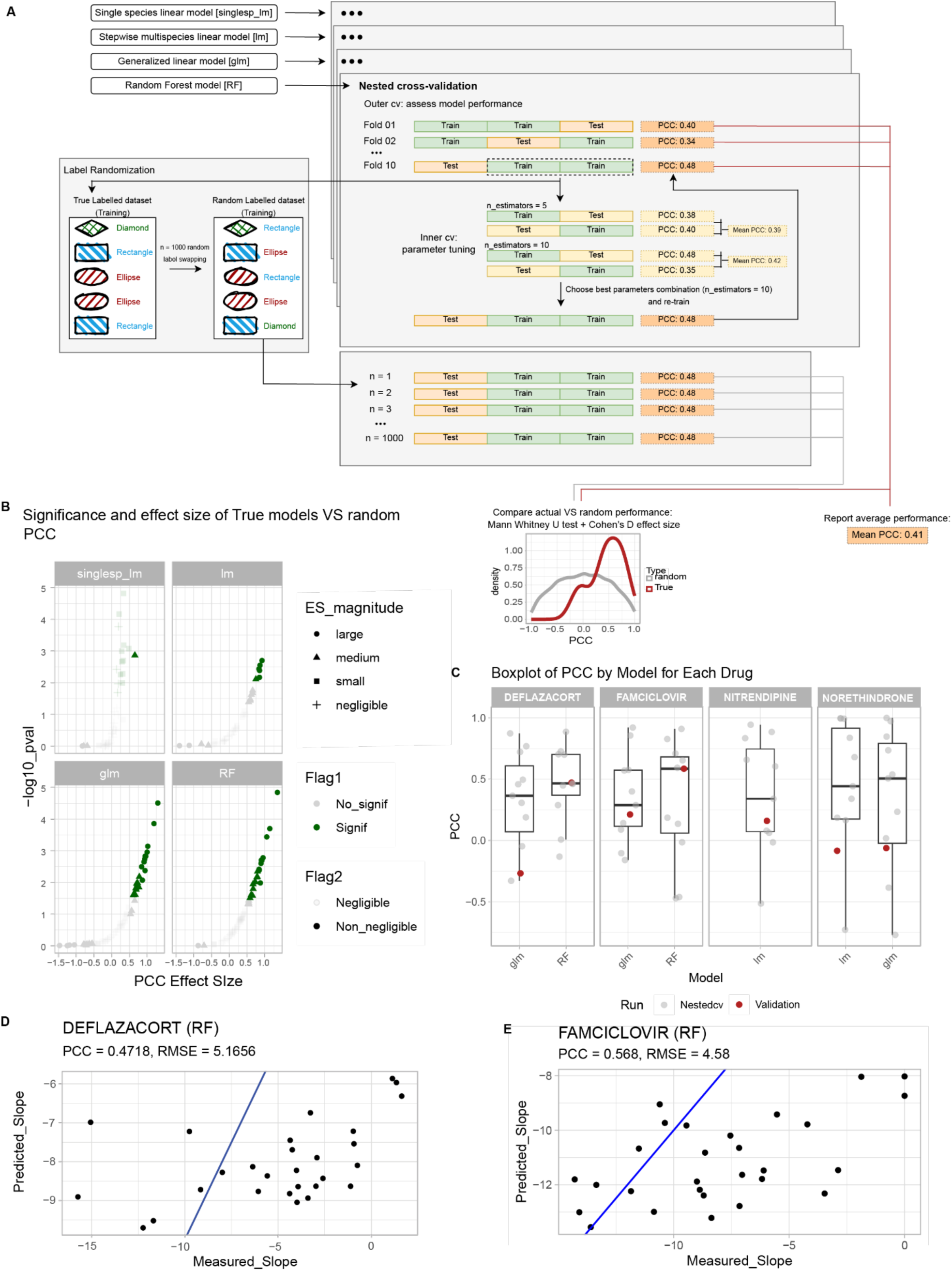
Overview of modelling results. (A) Schematic representation of the modelling approach, including single species linear models, stepwise multispecies linear models, generalized linear models and random forest, trained and tested by nested cross-validation. To assess whether model performances were better than random, they were compared to the model predictions of 1000 random models built with the same set of features following random swapping of the outcome vector labels. (B) True and random model performance (as measured by PCC) were compared by assessing (i) significantly better performance, measured as one-tailed Mann-Whitney U test and (ii) non-negligible effect size, measured via Cohen’s d. Significant and non-negligible results are highlighted in non-transparent colors. (C) Overall comparison of the performance obtained when validating the transferable models on a previously unseen set of 28 human gut communities: boxplot representing the distribution of PCC over the 10 nested-cv folds used to train the models (also reported as gray dots) and the corresponding PCC obtained on the validation data (in red). (D-E) Scatterplot of measured vs. predicted biotransformation rates when the two best performing rf models are validated: results for the drugs deflazacort (D) and famciclovir (E), with RMSE and PCC indicated. Blue line represents the bisector, with points lying on the diagonal if predicted and measured biotransformation rates are the same.

In most of the combined linear models (31/50), glm (48/65) or random forest models (65/65), the model selected more than one predictor, suggesting that the combination of microbes predicts the observed biotransformation rates better than any individual species (Table S19). This is likely due to the fact that multiple species can contribute to the biotransformation of a given drug, both within and across communities. As a consequence, the complex modelling approaches resulted in superior overall model prediction accuracy with (i) PCC range increasing from [-0.598; 0.585] for single species linear models, to [-0.689; 0.491] for stepwise linear models, to [-0.719; 0.699] for glm and [-0.465; 0.726] for random forest; (ii) the average PCC between prediction and measured biotransformation rate of 0.0694 and 0.184 for glm and random forest models, respectively (Figure S6A), (iii) random forest performing significantly better than all other approaches (as assessed via ANOVA with fdr adjusted p-values for multiple pairwise comparisons, Table S20). Furthermore, the more complex modelling approaches also enabled predictions for drugs without prior information on gut microbial biotransformation (n = 8 drugs) or in the absence of experimentally validated drug-metabolizing strains in the communities (n = 5) (Table S21), two of which with moderate correlation with the observed biotransformation rate (PCC = 0.521, 0.489 for prednisone and benzobromarone, respectively).

To further assess the quality of model predictions we verified that: (i) the prediction was better than random, by comparing it to the prediction obtained from 1000 random models built over the same cohort by randomly swapping the outcome vector and (ii) the prediction was better than baseline by assigning to all communities the average biotransformation rate observed in the cohort (Figure S6B-C). 49 models for a total of 32 drugs showed a significant (one tailed Mann Whitney U test fdr-corrected p-value <=0.1) and medium to large (as measured with Cohen’s d effect size) improvement in PCC compared to random models (Figure 4B). These 49 models contain 1 single species linear model, 6 stepwise linear models (4 of which with better than baseline performance), 21 glm and 21 RF models (Table S22). Taken together, these analyses suggested that gut microbiota drug biotransformation rates can be predicted from microbiome composition for 32 of the 66 drugs that show the strongest interpersonal differences in gut microbiota drug metabolism.

To validate the predictive models, we next examined an independent drug biotransformation data set available for deflazacort, famciclovir, nitrendipine and norethindrone acetate incubated with 28 human gut microbial communities from a distinct cohort of human volunteers whose samples were collected and stored using different methods^4^. Average performance of the transferable models (Figure S6D) for the four drugs are compared in Figure 4C and Table S23 both for the nested cross validation and for the external validation set. When applied to the taxonomic composition of the 28 microbial communities in the validation set, PCC between predicted and observed biotransformation rate was highest for both deflazacort and famciclovir when using the random forest model predictions, reaching 0.47 and 0.57, respectively (Figure 4D-E).

Together, these results indicate that the models trained on the data collected in this study exhibit moderate (and significantly better than random) performance when applied to community composition data from independent datasets. Additionally, the fact that the random forest models performed best on the validation data set further emphasizes that multi-taxa models are more robust and are hence most suitable for transfers between datasets.

### Interpersonal differences in gut microbial drug metabolism in preclinical animal models

Human drugs are typically tested in preclinical animal models prior to human trials; approximately 70% of candidate drugs are sidelined at this stage in the drug development process^29^. To test the extent to which microbiota drug metabolism activities observed in human microbiomes are recapitulated in microbiomes of preclinical animal models, we next collected 29 gut microbiome samples from six preclinical model species (mouse (n=8), rat (n=4), rabbit (n=4), dog (n=3), pig (n=4), monkey (n=6); Table S24) under the same collection methods used for the human-derived samples. As above, we measured the metagenomic composition and biotransformation kinetics of the 271 medical drugs by each of these communities, which we designate as “animal microbiome samples” for simplicity.

Differences between human and non-human gut microbiome composition have been ascribed to differences in diet, host anatomy and physiology^30–33^. Beta diversity analysis of the collected preclinical animal microbiome samples revealed (i) limited overlap between human and animal samples, independent of the animal species (Figure 5A), (ii) reduced diversity across individual preclinical animals from the same species compared to human microbiome diversity (as tested by betadisper test, Table S25), and (iii) similarity between rodent samples (mouse and rats). Despite these expected differences in microbiome composition, we found 197 of the 370 bacterial species observed in at least one of the 60 human gut communities were also present in the gut microbiomes of at least one of the tested preclinical animals. These 197 bacterial species cover on average 84.4% of the overall abundance of human samples and 93.5% of the animal samples (Table S26), and thus suggested potential transferability of the human gut microbial drug biotransformation capacity to the animal models.

**Figure 5.**
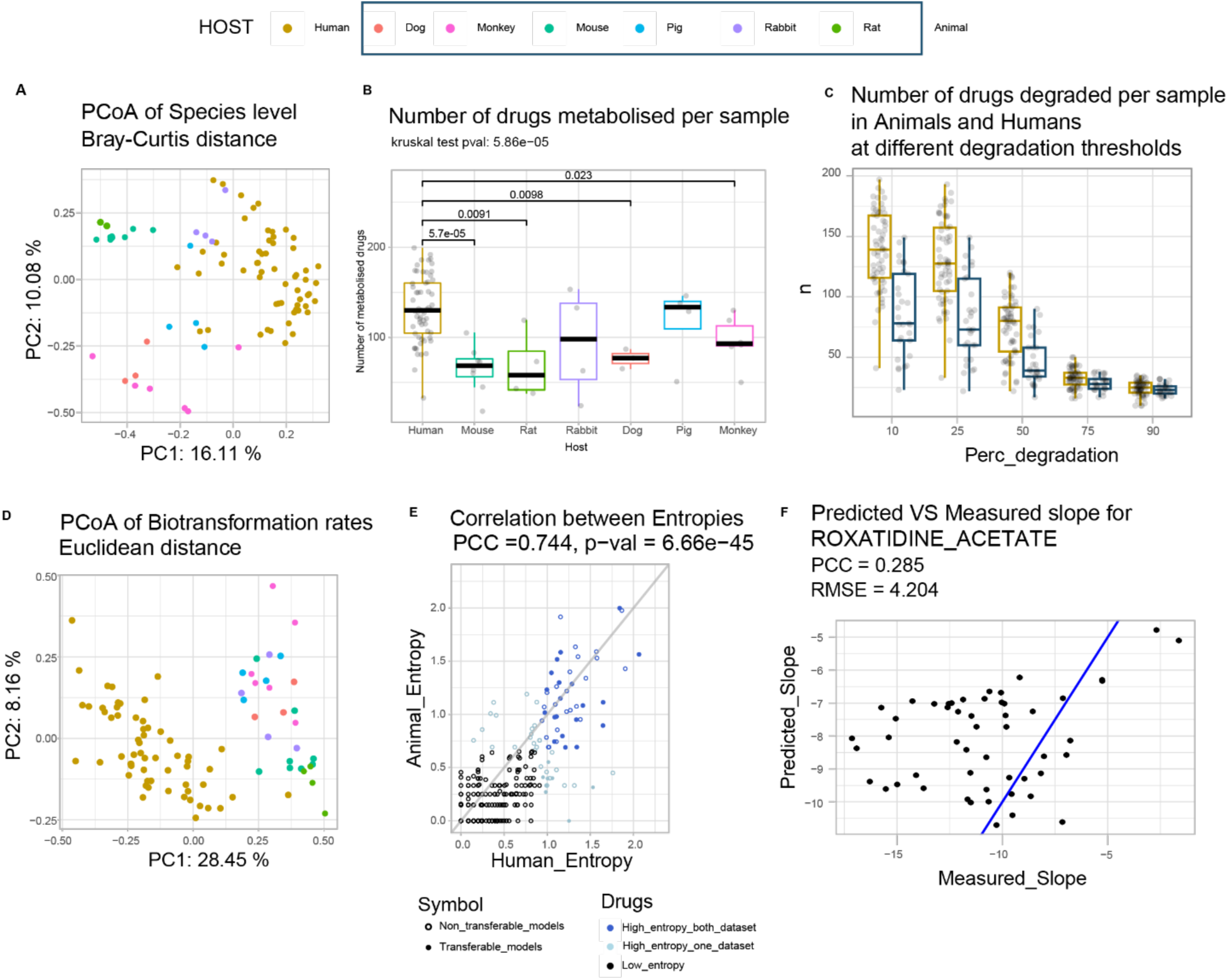
Similarities and differences between human and animal model gut communities and their drug biotransformation. (A) PCoA of Bray-Curtis dissimilarity between gut microbiota samples from human and preclinical animal models. (B) The number of metabolized drugs (25% reduction in parent compound levels) across microbiome samples from humans and preclinical animal model species. (C) The number of metabolized drugs in human and animal model samples across different thresholds for reduction of parent compound levels; across most thresholds, the animal preclinical models biotransform fewer drugs compared to the human cohort. (D) PCoA of Euclidean distance based on biotransformation rates measured in the gut microbiota of human and preclinical animal models. (E) Correlation between entropies calculated from the biotransformation rates for each drug in human (x-axis) and animal (y-axis) samples. Most (44/66) of the high entropy drugs identified in the human cohort are also exhibit high entropy in the animal cohort (dark blue circles). Among them, 24 transferrable models perform better than control models (random outcome vector) in predicting human biotransformation rate from models trained on animal data. (F) Performances are on average moderate or poor for all models trained on animal microbiome composition/drug metabolism data and then applied to human microbiome composition data to predict human microbiome drug metabolism. However, RF shows consistently the highest performance. Here, the best model is reported (drug roxatidine acetate), with a correlation between predicted and observed biotransformation rates equal to 0.285.

To identify drugs metabolized by the gut microbiota of the preclinical animal models, we compared drug levels before and after the biotransformation assays as described above (Table S27). While 8 drugs could not be quantified, we found that 223 of the remaining 263 drugs exhibited at least 25% reduction in parent compound levels by at least one animal-derived gut community (adjusted p-value <0.1, see Materials and Methods). The 29 animal microbial communities biotransformed between 24 and 143 drugs with an average of 85 drugs (Table S28). Twelve drugs could be biotransformed by all animal-derived microbial communities (*i.e.*, biperiden, bisacodyl, bromocriptine mesylate, cyproterone acetate, diacetamate, duloxetine, fluvoxamine maleate, melphalan, nitrendipine, oxethazaine, phenazopyridine and tinidazole). Two of these 12 drugs (bisacodyl and diacetamate) were also biotransformed by all human gut communities and they have previously been described to be hydrolysed by a diverse set of human gut bacteria^4^. Notably, approximately two-thirds (27/42) of the drugs that were not biotransformed by any of the tested animal-derived communities were significantly metabolized by at least one of the human samples. Comparison of the number of biotransformed drugs between host species revealed that the rat microbiome samples show the smallest (average = 38) and the pig samples the highest (average = 130) number of metabolized drugs (Kruskal-Wallis test, p-value = 6.50e-3). Moreover, the number of drugs biotransformed per sample is significantly lower in animal compared to human-derived gut communities, either when all animal species are collapsed or analysed individually (Kruskal-Wallis test, p-value = 3.25e-07 and p-value = 5.86e-05, respectively) (Figure 5B). To test how the observed difference in drug biotransformation capacity between the human and animal microbiota depends on the biotransformation threshold, we compared the number of metabolized drugs between human and animals at different biotransformation cut-offs (*i.e.*, 10, 25, 50, 75, and 90% reduction in parent drug levels). We found that the number of drugs biotransformed per sample is significantly different between animal-and human-derived gut communities for all biotransformation thresholds between 10%-75% (Figure 5C, Table S29), with 75% showing the smallest effect size (eta squared based on the H-statistic, eta = 0.0517). To determine whether human and animal samples efficiently (>75% reduction in parent drug levels) target the same drugs, we identified 99 drugs that exhibited >75% biotransformation in at least one human or animal sample. Only 19 of these 99 drugs passed this threshold for all animal and human samples, and approximately one-third (32/99) passed the 75% biotransformation cutoff for all human samples but not for any animal sample. (Figure S7A).

We next calculated the drug biotransformation rates for all animal samples and all drugs. Consistent with microbiome composition similarities, clustering the samples based on biotransformation rates groups the animal samples separately from human samples while grouping rodent samples together (Figure 5D). Despite the described differences between gut microbial drug biotransformation rates between human and animals, we found comparable biotransformation rates for 19% of drugs (*i.e.*, distribution of biotransformation rates not significantly different between the human and animal cohort, Kolmogorov-Smirnov test for distributional equivalence, fdr corrected p-value >0.05) (Table S30). For example, the calculated rates of ethynodiol acetate biotransformation ranged from-0.183 (equivalent of-0.366 µM/h) to 0 in human microbiome samples and from-0.182 to 0 across preclinical animal samples (Figure S7B). However, for the great majority (81%) of drugs, the biotransformation rates measured across the preclinical animals fail to capture the overall diversity of the human microbiota biotransformation rates (*e.g.*, verapamil, Figure S7C), underestimate human microbiota drug metabolism (*e.g.*, sulfinpyrazone, Figure S7D) or fail altogether to identify drug biotransformations that occur in the human-derived microbiota samples.

To test how well preclinical animal models reflect the observed interpersonal differences in human gut microbial drug biotransformation for specific drugs, we compared biotransformation entropies for each drug between the human and animal measurements. Although we found a significant correlation between the entropies measured in the human and animal cohorts (PCC=0.744, p-val=6.66e-45), only 44 of the 66 high-entropy drugs identified in the human cohort also exhibited high entropy in the animals (Fig 5E). For 20 of these 44 drugs, we had a predictive model that was successfully trained and cross-validated on the human cohort data and that we could test for their potential transferability to preclinical animal models (Table S22, Fig 5E, highlighted as filled circles). For this subset of 20 drugs, we conducted an explorative analysis to test whether human gut microbiota drug metabolism could be predicted based on the data from preclinical animal models. To this end, we re-trained the 32 transferable models for these 20 drugs using the composition and the measured biotransformation rates of the animal-derived gut communities to then use these models to predict human microbiota drug biotransformation rates. Out of these, one drug failed to be modelled from animal samples; the two drugs that could only be modelled with linear model failed to predict biotransformation rate variability in the human data (*i.e.*, they gave the same prediction for every sample due to a lack of overlap of selected microbial species between animals and human). Of the remaining 29 models, 24 (14 RF models and 10 glm models) were able to predict biotransformation rates, while 5 glm models were non-significant (Table S31) (Fig 5F showcasing a good prediction case). Independently from the number of predictors retained by the models, performances were moderate to poor, with average PCC between predicted and observed biotransformation rate equal to 0.107 and 0.0530 for RF and glm, respectively. Nevertheless, performance of RF models significantly correlated with the estimated average performance observed in humans (Spearman rho = 0.556, p-value = 0.0420). These exploratory results highlight the potential limitations of preclinical animal models modelling microbiota drug biotransformation, both in terms of overall prevalence and of interindividual variability.

## Discussion

A central challenge in studying gut microbial drug biotransformation is the complexity caused by both the natural diversity of gut microbial communities and the wide chemical diversity of drug molecules.^34^ In this study, we combined metabolomics and metagenomics analyses to systematically assess the occurrence and inter-individual variation of biotransformation of 271 drugs by 89 gut microbial communities derived from humans and common preclinical animal models, and provide this data as a resource for future study. This resource includes more than 868,000 measurements that capture 24,119 drug-microbiota interactions over 9 timepoints, each in quadruplicate, plus negative controls at multiple pH levels for each drug as well as metagenomic data for each microbiome sample.

This dataset reveals several key findings. First, more than 90% of the tested drugs were metabolized by at least 25% by at least one of the tested communities highlighting the diverse repertoire of drug-metabolizing enzymes encoded in gut microbiomes. We identified several drugs that had not (to our knowledge) been previously reported to be metabolized by individual gut bacterial species, supporting the hypothesis that microbial drug biotransformation includes activities that only emerge in a community context^8,19^.

Second, the dataset generated in this study provides an opportunity to characterize inter-individual and between-drug variability in drug biotransformation rates. Human gut microbiomes from individual donors metabolized as few as 35 and as many as 206 of the tested drugs, underscoring significant interpersonal differences. Considering the relatively small size of the tested human cohort (all identified as westernized), it is likely that interpersonal differences in gut microbiota drug biotransformation are even more pronounced across the general population^34^. While many drugs exhibited high entropy (interpersonal variation in biotransformation rate), we observed that certain sets of high-entropy drugs were metabolized rapidly or slowly by the same microbiome samples. In some cases, these drugs shared overall chemical similarity (as in the case of palperidone and risperidone) or the presence of specific functional groups (*e.g.*, famciclovir, roxatidine acetate, and deflazacort). However, the presence of a given functional group alone seems generally insufficient to explain gut microbial biotransformation rates of different drugs (*e.g.*, phenazopyridine and sulfasalazine share an azo group, but their biotransformation rates across the human microbiome samples do not correlate significantly, PCC = 0.076, p-val= 0.565).

Third, this data resource facilitates development and testing of modelling approaches of increasing complexity for predicting the observed interindividual variability in drug biotransformation rates from community composition data. Simple linear correlation analysis exemplified how drug biotransformation data of individual strains can explain microbiota community metabolism for certain drugs. However, for most of the drugs investigated, the observed relationship between community composition and biotransformation appeared to involve more complex behavior. To address this complexity, we employed a multi-model predictive strategy starting with the simplest model possible (linear models) based on prior knowledge of bacterial biotransformation. We then progressively incorporated additional complexity factors, including overall community composition, enzymatic profiles, and potential non-linearity in the relationship between predictors and outcomes. Although our modelling approach is limited (*e.g.*, with respect to the models included and dataset diversity and size) and microbiome metabolism is too diverse for all putatively relevant predictors to be captured, we were able to predict gut microbial biotransformation rates with mid-to-high correlation with the actual measured rates for certain drugs.

Fourth, this dataset enables direct comparison of gut microbial drug biotransformation between humans and commonly used preclinical animal models. Notably, no single preclinical model adequately captures the breadth or diversity of drug biotransformation observed in the human gut microbiota, even after accounting for differences in sample sizes across groups. In our experimental setup, only the combined data from all seven preclinical models was able to recapitulate a portion of the inter-individual variability seen in human gut microbial drug metabolism. Overall, our analysis underscores several limitations in predicting human microbiome-drug interactions based on preclinical animal microbiotas: i) We identified several drugs, including celecoxib and nefazodone, that displayed substantial interpersonal variability in human microbiota biotransformation but were not biotransformed by any non-human samples. ii) Only 66% (44/66) of the drugs that showed the greatest inter-individual variation in humans exhibited similarly high variability in the animal cohort. iii) Restricting analyses to a single animal species substantially increased the number of drugs that were biotransformed in human samples but not in the tested animal model. For example, 32 drugs were biotransformed in >75% of human samples but no animal samples, and restricting analysis to a single animal species adds 12 (pigs) to 32 (rats) additional drugs that were missed in the animal model. These findings, together with the relatively high overlap in detected microbial species across models (53.24%), suggest that rare or low-abundance human gut microbes may play important roles in mediating microbiome-driven drug biotransformation.

Our study has several limitations. The relatively small sample size (both human and animal) reduces the generalizability of our conclusions and the robustness of the models, and the high-throughput nature of this approach precludes incorporation of host ADME in these measurements. Thus, we present our approach as a proof of concept for modelling microbiota-mediated biotransformation rates, rather than as ready-to-use models. Larger and more diverse cohorts, including individuals of different ethnicities, sexes, and lifestyles, will improve generalizability. Our choice of modelling methods was constrained by sample size, prior knowledge, and the need for flexible approaches, but may not represent the most efficient strategy. Furthermore, low-prevalence or low-abundance taxa were excluded to reduce feature space, though such taxa may still influence drug biotransformation. Finally, our models explore a link between taxa/gene abundance and biotransformation rate. While our results support this association in some cases, we expect that other factors (*e.g.*, community interactions, drug bioaccumulation, gene regulation) will also play critical roles in determining drug metabolism and contribute to the modest predictive power observed in models based on taxa and gene abundance alone.

Overall, our results demonstrate how high-level descriptions of chemical and ecological diversity are generally insufficient to capture drug biotransformation in the community context, and that smaller scale features (*e.g.*, chemical or community features) likely need to be investigated on a per-drug basis to better predict interindividual variation in microbial drug metabolism. To this end, our dataset provides a resource for hypothesis generation by highlighting several potential features that contribute to biotransformation, as well as providing a testing platform for modelling approaches. In our limited attempt at modelling biotransformation rates, we showcase both the challenges and opportunities posed by the combination of community composition and its functional reservoir in predicting community drug biotransformation.

## Acknowledgments

We thank the Zimmermann and the Goodman labs for helpful discussions, G. Barnett and the Yale Animal Resource Center for providing the non-human fecal samples tested in this study, T. Wu and the Yale Center for Molecular Discovery for technical assistance, and B. Zhang and S. Devendran for help with manual inspection of mass spectrometry data. This work was supported by NIH grants R01AT010014, R35GM118159, and R01DK13379 to A.L.G, P30AG017266 and 5R01AG060737-01 to P.H. and the European Molecular Biology Laboratory, Daimler-Benz Fellowship and an ERC Starting Grant (GutTransForm ID 101078353) to M.Z. EMBL IT Support is acknowledged for provision of computer and data storage servers.

## Author contributions

M.Z. and A.L.G. conceived and initiated the project; P.H. and F.E.R. provided human faecal samples from the Wisconsin Longitudinal Study; M.Z. performed the experiments; E.M. analysed the data, performed statistical analyses and prepared graphical illustrations. E.M. and M.Z. wrote the initial draft manuscript; All authors reviewed and edited the manuscript.

## Data and code availability

Raw sequencing data have been deposited on the ENA server, with accession number PRJEB37062.

All raw metabolomics data and correlated metadata are deposited in Metabolights under study number MTBLS6166. All other data are available in the main text or the supplementary materials, and additionally available on Zenodo with DOI 10.5281/zenodo.13943903. Moreover, all community drug biotransformation data were also summarized and publicly shared in Chembl v.36. Analysis scripts are available on GitHub (https://github.com/ZimmermannLab/Community_drug_metabolism.git)

## Tables with titles and legends

**Table S1:** Drugs used in this study, name, formula, mass, logP, chromatographic retention time and additional relevant chemical metadata.

**Table S2:** Community composition (as assessed via metagenomic shotgun sequencing) of 60 human gut communities used in this study.

**Table S3:** Drug screen results: parent drug quantification (via LC-MS) for 271 drugs in 60 human gut communities plus non-bacterial controls over 9 timepoints in quadruplicate

**Table S4:** Number of drugs identified as metabolised by each community at different biodegradation tresholds (0.9, 0.75, 0.5, 0.25, 0.1) and their respective cumulative sums.

**Table S5:** Detail of number of communities degrading a drug at least 25%, and type of identified metabolism (quick or normal).

**Table S6:** Biotransformation rate estimated for each drug and sample (microbial community).

**Table S7:** Estimated Shannon entropy of biotransformation rates for each drug.

**Table S8:** Tanimoto distance matrix among tested drugs

**Table S9**: Over representation analysis result of chemical groups of the drugs according to their interpersonal variation (i.e., entropy) in gut microbiota biotransformation rates.

**Table S10:** Pearson correlation of biotransformation rates for the drugs forming a network of six distinct clusters in Figure 2B.

**Table S11:** Reference protein sequence of 31 gut bacterial genes that have previously been experimentally validated to biotransform 21 of the tested drugs

**Table S12:** Drug degraded by each of the 31 genes listed in Table_S11

**Table S13:** Quantification of each of the 31 reference protein marker abundance listed in Table_S11 in 60 human metagenomes

**Table S14:** Spearman correlation between estimated biotransformation rate of each drugs and the abundance of the associated metabolic gene(s).

**Table S15:** Spearman correlation between estimated biotransformation rate of each drugs and the abundance of each taxonomic feature across all communities (8 phyla, 14 classes, 19 orders, 43 families 108 genera and 370 species).

**Table S16:** Identification of 68 strains previously assayed in axenic cultures for the biotransformation of the 66 drugs

**Table S17:** Number of individual strains potentially metabolising ach of the 66 drugs identified in the communities

**Table S18:** Average Pearson Correlation Coefficient (PCC) between observed and predicted biotransformation rate measured over 10 nested cross-validation for each of the four models developed for each of the 65 drugs (single species linear model, stepwise linear model, generalized linear model, random forest). For the linear models, significance is reported as well.

**Table S19:** Optimal number of variables retained for each drug with each of the modelling approaches.

**Table S20:** Analysis of variance result for comparison of PCC performances across the four different modelling approaches.

**Table S21:** Drugs that could only be modelled with glm or RF approach, for absence of previous evidence or the impossibility to quantify the responsible species in the communities

**Table S22:** Models with performance significantly better than random or baseline models **Table S23**: Comparison of performances over nestecv and on actual validation set for 4 transferable models

**Table S24**: Metadata of collected animal samples

**Table S25**: Beta dispersion test result and Tuckey HSD for differences in dispersion observed when using Bray-Curtis distance to assess microbiota composition differences between human and animals.

**Table S26**: Abundance of 197 bacterial species found in common between human and preclinical animal communities

**Table S27**: Drug screen results: parent drug quantification (via LC-MS) for 271 drugs, 29 animal gut communities plus non-bacterial controls over 9 timepoints in quadruplicate

**Table S28**: Number of drugs identified as metabolised by each animal community

**Table S29**: The difference in number of drugs being identified as metabolised by animal and humans does not depend on the chosen biotransformation threshold

**Table S30:** For 19.5% of drugs comparable biotransformation rates could be observed between the human and animal cohort, as tested by Kolmogorov-Smirnov test for distributional equivalence.

**Table S31:** Performance of transferable models trained on animal communites and tested on human communities

## Methods

### Chemicals

Drugs were picked from the Pharmakon1600 library (MicroSource Discovery Systems) as 10mM stock solutions and combinatorial compound pooling was performed by the Yale Center for Molecular Discovery as previously described^1^. LC–MS-grade solvents were purchased from Fisher Scientific.

### Human and preclinical animal model microbiome samples

Fecal samples were previously collected using the fecal aliquot straw technique^2^ and stored at-80°C as part of the Wisconsin Longitudinal Study (WLS)^3^. WLS specimen collection was approved by the UW-Madison Internal Review Board (2014-1066, 2015-0955) and written informed consent was obtained in the original study^3^. Each sample was de-identified by WLS staff before involvement in this study; sample numbers used in this study are not associated with study volunteer names or other identifying information. No metadata was received about the subjects. Fresh fecal material from animals housed at the Yale Animal Resource Center (YARC) was collected and cryopreserved using the same protocol as for the human fecal samples.

### Drug biotransformation assays

Drug biotransformation assays were performed inside a flexible anaerobic chamber (Coy Laboratory Products) containing 20% CO_2_, 10% H2 and 70% N_2_. 5 mL of gut microbiota medium (GMM)^4^ was inoculated with 100 mg of frozen fecal material and incubated at 37°C for 12 to 16 h. Drug-conversion assays were performed as previously described^1^. In brief, bacterial cultures were diluted into fresh, pre-reduced GMM (5-fold diluted in MilliQ water to a final volume of 250 µL) containing the tested drugs at 2 μM final concentration of each individual compound in the combinatorial pools. Biotransformation assays were performed for 24 hours at 37°C under anaerobic conditions and 20 µL of whole-culture samples were collected at 0,1, 2, 4, 6, 8, 12, 18, and 24 h of incubation, flash-frozen, and stored at−80 °C until further processing for analysis by LC–MS.

### DNA extraction and sequencing

DNA of bacterial community samples was extracted from a biomass pellet of 500-μl community cultures before the biotransformation assay as previously described^5^. Library preparation and sequencing were performed at the Yale Center for Genome Analysis. The Kapa Biosystems HyperPrep kit was used for the preparation of the metagenomic sequencing libraries. The 2×150-bp sequencing was performed on an Illumina Novaseq 6000 instrument with a S4 flow-cell to a target depth of ∼20 Mio reads per sample as previously described^1^.

### Metagenomic data analysis

Metagenomic sequencing data preprocessing and analysis was performed with the bioBakery tools suite^6^. Paired-end reads from each sample were quality checked using FastQC v0.11.5 (available online at: http://www.bioinformatics.babraham.ac.uk/projects/fastqc/) and filtered for host DNA contamination and adapter trimmed with KneadData v0.7.2 (using trimmomatic-0.33^7^ and bowtie v. 2.3.5^8^). The reference database for each of the host (human and preclinical animal models) was created using the references provided in [Table S32] (all reference sequences have been downloaded from Ensembl release 98 on 20-nov-2019). An overview of QC and host filtering data is reported in [Table S33]. Filtered paired-end reads were merged before further processing. All downstream analyses were performed on pre-processed reads unless otherwise stated.

Taxonomic assignment was performed with MetaPhlAn v2.7.7^9^ on the filtered and merged sequencing data. All outputted tables were then merged into a summary taxonomic table with the merge_metaphlan_tables function (Table S34, Table S35).

The taxonomic tree representing bacterial species identified was visualized with GraPhlAn v1.1.3^10^. Protein sequence identification and abundance quantification was performed using ShortBRED v.0.9.5^11^. Target protein sequences, previously reported in Zimmermann et al.^1^ and Baghai Arassi et al.^12^, were retrieved from NCBI Batch Entrez^13^ and are listed, together with their amino acid reference sequence, in [Table S11]. ShortBRED markers were created with shortbred-identify function using Uniref90 as a reference^14^ (downloaded on March 5th, 2020, from ftp://ftp.uniprot.org/pub/databases/uniprot/uniref/uniref90/). Protein marker abundance was quantified using the created reference markers with shortbred_quantify function. All resulting tables were joined into a summary quantification table (Table S13, Table S36).

### Statistical analysis of metagenomic data

Data processing, statistical analysis and plotting were performed in R version 4.2.0 [R Core Team (2022). R: A language and environment for statistical computing. R Foundation for Statistical Computing, Vienna, Austria. URL https://www.R-project.org/]. All taxonomic tables were imported, and the following criteria were used to filter ASV tables: for samples, only those with (i) final proportion of retained reads (after trimming and host decontamination) >90% and (ii) percentage of reads taxonomically annotated to bacteria kingdom >75% were retained. For features, only those (i) minimum prevalence of 2 samples (*i.e.*, singletons, features found in one sample only, were excluded). One sample was excluded because it failed quality check (percentage of reads taxonomically annotated to bacteria <75%).

Metagenomes of 2803 healthy Western donors were obtained were obtained using the R package curatedMetagenomicData, with the “*metaphlan_bugs_list.stool” option to download all metaphlan tables available for stool samples; downloaded samples were then filtered for: (i) body_site equal to ‘stool’, (ii) disease metadata equal to ‘healthy’, (iii) age_category equal to ‘adult’ (iv) study_condition equal to ‘control’ (v) non_westernized equal to ‘no’ (i.e., westernized only were selected), (vi) country to be not equal to any Asian/African country. The resulting table with ASV relative abundance is reported in Table S37 and the associated metadata in Table S38.

Beta diversity analysis, Permutational Multivariate Analysis of Variance (PERMANOVA)^15^ and Multivariate homogeneity of groups dispersions (betadisper)^16,17^ were performed using the vegan package, using the vegdist, adonis2 and permdisp function, respectively. PERMANOVA analysis was performed with block design to consider the presence of different studies. First, the test was conducted on the whole dataset, with 2999 permutations; then, to mitigate the unbalanced comparison between our dataset and the reference set, 1000 bootstrap permutations of the test were performed, where each time 30 random samples were extracted from our 60 samples, and 30 random samples were extracted from the 2803 deposited samples; there, the permanova test was run with 999 permutations each.

Alpha diversity analysis was performed using the DiversitySeq package with the Chao1 index^18^. Stacked barplot of species (as well as genus, family, order, class, phylum) were obtained by capping the minimum represented relative abundance to 1%. Sample hierarchical clustering based on community species relative abundance or on the relative abundance of 63 representative species was obtained using Bray-Curtis dissimilarity^19^ and average linkage. Heatmaps were visualized using the pheatmap package (https://CRAN.R-project.org/package=pheatmap). Tanglegram construction, dendrogram comparison, difference, entanglement and correlation (measured as cophenetic distance) were obtained using the dendextend package.

Mantel test of Bray-Curtis beta diversity and Euclidean based drug biotransformation diversity was computed using the mantel function from the vegan package, using method pearson and 999 permutations. Individual species with known biotransformation activity against individual drugs were derived from Zimmermann et al.^1^, and quantified in the communities by exact match of species name.

### Targeted Metabolomics analysis

#### Sample preparation and LC-MS analysis

Samples were prepared for LC–MS analysis by organic solvent extraction (acetonitrile:methanol, 1:1) at −20°C after the addition of internal standard mix (sulfamethoxazole, caffeine, ipriflavone and yohimbine each to a final concentration of 80 nM) as previously described^1^ using an EpMotion liquid handling system (Eppendorf). LC-MS analyses was performed as previously described^20^. In brief, chromatographic separation was performed by reversed-phase chromatography (C18 Kinetex Evo column, 100 mm × 2.1 mm, 1.7-mm particle size, and according guard columns, Phenomenex) using an Agilent 1200 Infinity UHPLC system and mobile phase A (H_2_O, 0.1% formic acid) and B (methanol, 0.1% formic acid), and the column compartment was kept at 45°C. Five µL of sample were injected at 100% A and 0.4 ml/min flow, followed by a linear gradient to 95% B over 5.5 min and 0.4 ml/min. To ensure reproducible chromatographic separation (retention shifts between samples <2% or 0.15 min) columns were changed after 1,000 sample injections. The qTOF instrument (Agilent 6550) was operated in positive scanning mode (50–1,000 *m*/*z*) with the following source parameters: VCap, 3,500 V; nozzle voltage, 2,000 V; gas temperature, 225 °C; drying gas 13 l/min; nebulizer, 20 psig; sheath gas temperature 225 °C; sheath gas flow 12 l/min. Online mass calibration was performed using a second ionization source and a constant flow (5 μl/min) of reference solution (121.0509 and 922.0098 *m*/*z*). The MassHunter Quantitative Analysis Software (Agilent, version 10.0) was used for peak integration based on retention time and accurate mass measurement of chemical standards.

#### Quality Check

Targeted data for each drug and community was imported in R version 4.2.0 [R Core Team (2022). R: A language and environment for statistical computing. R Foundation for Statistical Computing, Vienna, Austria. URL https://www.R-project.org/] for further data processing, statistical analysis and plotting, by relying on the data.table package for efficient data manipulation. Both RT drift and plate border effect were checked, and no sample was identified as outlier (and therefore had to be excluded) according to such criteria. Complete raw metabolomic datasets are reported in Table S3 and Table S27 for the human and animal cohort respectively.

To reduce the overall level of noise, a noise threshold of 5000 counts AUC was set and the following filtering criteria were applied: (i) all quantified drugs with an intensity below the noise threshold, were set as equal to the noise threshold itself (excluded data points are detailed in Table S39); (ii) all quantified drugs with an intensity below 1% of the drug average intensity in the no-bacterial control samples, were set as equal to the noise threshold (excluded data points are detailed in Table S40); (ii) all kinetic profiles including less than 3 timepoints were excluded. Internal standard normalization was applied as reported previously^1^: yohimbine, caffeine and sufamethoxazole were used as internal standards and their median fold change per sample and timepoint was used to correct measured area.

Outliers were identified both per time course and timepoint as follows: (i) as values outside trimmed mean ± 2 trimmed standard deviations, with trimming proportion set to 0.2 (so two outliers per time course could be present without altering the mean/sd); (ii) as timepoints whose standard deviation of the 4 replicated pools lies outside the overall timepoints standard deviation mean ± 2 trimmed standard deviations. Excluded data points are detailed in Table S41. Time course data for each drug and community were plotted and compared with raw data time course to check the efficacy of the quality check, normalization and outlier removal process.

Noisy drug/community profiles excluded from further analysis were identified as those time trends whose normalized intensity was varying for more than 2 consecutive timepoints for more than 5% of the value of the previous timepoint in opposite directions (*e.g.*, increasing/decreasing/increasing or decreasing/increasing/decreasing) for at least 2 samples and having a standard deviation of the log2 normalized area >1. Moreover, drug/community profiles showing a standard deviation of the log2 normalized area greater than 2 for at least 2 timepoints and for at least 2 samples were also excluded (excluded drug/community are detailed in Table S42). Lastly, drugs that in the previous steps lost data for more than 50% of the communities, or that lost all the no-bacterial control samples were excluded from downstream analysis.

Fold change (FC) was computed between each timepoint of a time course and timepoint 0. Significance was assessed with a Student t-test, as implemented in the t.test function, and its p-value corrected for multiple hypothesis testing with fdr correction^21^, as implemented in the p.adjust function. All FC computed are detailed in Table S43.

#### Drug biotransformation screen analysis

Each drug/community time trend was considered to be possibly describing a drug degradation event if it showed a FC significantly lower than a chosen FC cutoff for at least 2 consecutive timepoints and at least once among the timepoints 12h, 18h, 24h; fdr corrected p-values were deemed significant if below a p-value cutoff of 0.1. These criteria allowed us to divide drug/community time course analysis in two branches, described in more detail below.

Drug/community time trends not fulfilling the previous FC criteria could still describe a drug degradation event if the drug was already greatly degraded by the community before the first sampling point (hereafter referred to as ‘quick degradation’). To identify these cases, we compared time trends with a simulated trend for quick degradation, *i.e.*, a trend where each timepoint was represented by normally distributed samples with mean fixed to 5000 counts (*i.e.*, equal to the noise threshold) and standard deviation estimated from raw data before denoising (see “Targeted Metabolomics analysis: Quality Check”). Each timepoint of the drug/community time course was individually compared to the simulated trend with a one-sided Wilcoxon test, and p-values were then fdr corrected over the trend using the p.adjust function. If more than half of the timepoints in the trend were not distinguishable from the simulated quick degradation profile, but the drug could still reliably be measured in the no-bacteria controls, the drug/community time trend was labelled as a “quick degradation” event. Quick degradation drug/community are detailed in Table S44.

Drug/community time trends fulfilling the degradation FC criteria, were further compared to the corresponding no-bacterial control for the same drug, and were labelled as degrading events if either the no-bacterial control was showing no significant degradation over time or the community was showing a significantly higher degradation than the no-bacterial controls. To assess this condition, FC was computed between each timepoint of a time trend in communities and the corresponding timepoint in the no-bacterial control. Significance was assessed and fdr corrected as described above. As before, the drug/community time trend was considered to be describing a drug degradation event if it showed a FC significantly lower than the no-bacteria control for at least 2 consecutive timepoints. Degrading drug/community combination are detailed in Table S45.

To monitor the impact of the threshold choice on the number of communities identified as biotransforming each drug, we screened a vector of possible FC thresholds ranging from 0.1 to 0.9 (Figure S1B). Unless otherwise stated, we set our threshold to be 0.75, i.e. requiring a minimum of 25% reduction of the drug in the community. Moreover, to further account for variability in drug measurements, for each drug we calculated an adaptive fold-change threshold equal to the greater value between 25% or the mean of the fold changes, for which log2(fold change) > 0, plus 2 times the standard deviation^1^.

#### Quantification of drug biotransformation rates

All drug/community time courses were scaled to t0 (or to the t0 of the no-bacteria control for quick degradation events) and plotted with ggplot2. For each drug/community time course, initial slope was computed locally by feeding at least the first 4 timepoints to a linear model (Scaled_Area – Intercept ∼ 0 + Time, with Intercept fixed to 1), and iteratively adding more timepoints. The best model was chosen as the one showing the best adjusted R squared and its slope was returned as the initial slope for that drug/community and therefore as a proxy of drug degradation rate in that community. Diagnostic plots (*i.e.*, residuals, standardized residuals, residual versus fitted values and qqplots) were visually inspected for each time course (Fig S2).

Drug biotransformation rates were visualized as heatmap using the ggplot2 package, with non-negative slope values (possible due to measurement variability) forced to 0, and ordered according to Euclidean distance based hierarchical clustering of samples (Fig 1E).

Drug degradation rates of each community were ranked within drugs, and the Pearson correlation was measured with the function corr.test from the psych package.

Shannon Entropy^22^ was used to identify drugs showing high inter-community variability in drug degradation rate as computed by the entropy.empirical function of the package; results are reported in Table S7. The drugs showing the top 25% value for Entropy (66 drugs) were further analyzed by computing correlation among drug biotransformation rates for all communities using the Pearson Correlation Coefficient with the function corr.test from the psych package and visualized with the ggcorrplot package. Moderate to strong (PCC>=0.5) and significant (corrected p-value<0.05) correlated drugs were plotted with ggplot and visually inspected, and their R^2^ value was computed via linear modelling. Drugs showing a very high correlation in their biodegradation rate among the 60 samples (PCC ≥ 0.7) were further inspected for chemical similarity as described in the next paragraph.

#### Drug characterization

Chemical binary fingerprints and functional chemical group analysis were derived from Zimmermann et al.^1^. Chemical structure similarity for the drug clusters identified with hierarchical clustering was performed using the Structural Similarity Search on ChemMine^23^, which also provides the maximum common substructure between every pair of molecules selected. Tanimoto similarity between drugs was computed starting from drug SMILES codes (Table S1) using the ms.compute.sim.matrix function from the RxnSim package. Functional group enrichment depending on entropy of biotransformation rates observed among the 60 communities was computed with Fisher’s exact test as implemented in the fisher.test function, with both ‘greater’ and ‘lower’ option. P-values were corrected for multiple hypothesis testing using the fdr correction^21^, as implemented in the p.adjust function.

#### Metagenomics and Metabolomics data integration

Spearman correlations^24^ between all drug biotransformation rates and the relative abundance of all taxa at all taxonomic levels was computed with cor.test, and its significance was fdr corrected^21^, as implemented in the p.adjust function. All significant correlations are detailed in Table S15. Spearman correlation was also computed between 21 drugs and the quantification in each community of the corresponding protein encoding gene (or genes, when multiple were found in literature), reported to be responsible for drug biotransformation.

To test the possible influence of higher diversity on community drug biotransformation potential, Pearson correlation was computed between the total number of drugs degraded by each community and their corresponding alpha diversity measured as Chao1 index^18^.

We attempted to model gut microbiota drug biotransformation rate for the drugs showing the top 25% (n=66) value of Entropy using several approaches. Firstly, the drug degradation rate was modelled as a simple linear regression, using the *lm* function from the *stats* package, having as predictor the relative abundances of each of the species that have been previously demonstrated to be able to degrade the same drug in isolation^1^, i.e.

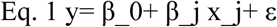

where y is the vector of degradation rates for each sample, 𝑥*_j_* is the relative abundance of species j in every community, 𝛽_0_ is the estimated intercept, 𝛽*_j_* is the estimated regression slope and 𝜀 is the error term.

Because our screen measured drug metabolism by bacteria in multi-species communities, we next implemented a stepwise approach to build a linear model accounting for all the species previously known to degrade the drug, and then, using a model stepwise selection based on the best Akaike information criterion (AIC)^25^, automatically retain the best combination of strains to build the final linear model. We implemented it using the *train* function with method=”lmStepAIC” and direction=”both” from the *caret* package, were the null model was assigned to the intercept only model, and the full model was assigned to the multiple linear regression having as predictors the relative abundance of all of the species that have been previously demonstrated to be able to degrade the same drug in isolation^1^, i.e.

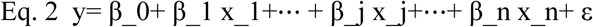

Whenever the number of predictors was higher than the number of samples available, therefore making the complete search unfeasible, the model was restricted to direction=’forward’.

To expand the modelling approach to species not previously investigated in isolation, but present in our communities, we also fit drug degradation velocity with Generalized Linear Models (GLMs) with penalized maximum likelihood, i.e. solving

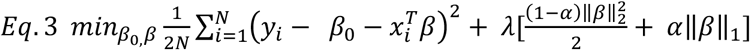

where 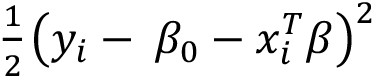 is the negative log-likelihood contribution of observation *i*, λ is the tuning parameter controlling the overall strength of the penalty, α is the elastic net penalty (tuned between 0.7 and 1, equivalent to the lasso regression), ‖𝛽‖_1_ is the *l_1_* norm of the beta coefficients and ‖𝛽‖_2_ is the *l_2_* norm of the beta coefficients. By automatically tuning both α and λ, GLM is able to (i) prevent overfitting and (ii) integrate feature selection in the algorithm itself.

Lastly, recognizing that this approach is still limited by the assumption of a linear relationship between taxa/gene abundance and drug degradation velocity, we also developed regression models based on Random Forest^26^.

Linear, stepwise and random forest model were trained using the function *train* from the *caret* package with method=‘lm’, method=‘lmStepAIC’ and method=‘ranger’ respectively; the glm model were trained using the *cva.glmnet* function from the *glmnetUtils* package. All prediction on the test set were performed using the *predict* function of the same packages.

To be able to assess model performance, the following information were retrieved for each of the chosen models: the number of tested predictors, the Root Mean Square Error (RMSE), the Pearson Correlation Coefficient (PCC) between measured degradation velocity and predicted velocity, the number n of optimal regressors used in the best model, and for linear models the significance of the chosen model (compared to the null model). All models’ performance was assessed in a nested cross validation (cv) setting (with 10 outer and 10 inner folds), which produce robust and unbiased performance estimates regardless of sample size^27^. To this aim, inner cv was used to optimize parameter tuning, and outer cv to assess model performance on previously unseen data.

All models used (i) as outcome variable the degradation rate of the drug in the corresponding community, estimated as described above, and (ii) as predictors the relative abundance of genes and taxa at all taxonomic levels as measured in each community, after Central Log Ratio normalization (clr)^28^, which has been shown to transform compositional data into symmetric and linearly related data^29^, which are therefore better suited for regression and modelling than total sum scaling only^30^. Moreover, taxa abundances were filtered to exclude (i) near-zero variance features, using the *nearZeroVar* function from the *caret* package (ii) low prevalence features, i.e. features observed in less than 10% of the training set, (iii) low abundance features, i.e. features whose maximum abundance across all training set sample was below 1%. To avoid information leakage, all filtering steps were performed only on training set samples and before applying normalization.

To assess the quality and significance of the prediction obtained from the models trained on our cohort we verified that: (i) the prediction was better than random, by comparing it to the prediction obtained from 1000 random models built over the same cohort by randomly swapping the outcome vector and (ii) the prediction was better than baseline, i.e. better than assigning to all communities the average biotransformation rate observed in the cohort. In order to guarantee reproducible results, the *set.seed* function was used before calling the *sample* function to randomly swap the labels of the training data. Performance comparison of true and random models was performed by computing (i) one sized Mann Whitney U test requiring true models to have significantly higher (fdr-corrected p-value <= 0.1) Pearson Correlation Coefficient with the hold out labels of the nested cross validation fold compared to random, as computed by the *wilcox.test* function and (ii) Cohen’s D effect size statistic as computed by the *cohen.d* function from the *effsize* package and requiring its magnitude to be equal to “medium” or “large” (i.e. |d|>=0.5).

When validation data were available either from other human or preclinical animal samples, the previously described methods for both taxonomic profiling of the metagenomic samples and estimation of drug degradation velocity were used, and then fed into the best performing model to assess its performance over the validation set.

## Supplementary Tables

**Table S3**: LC-MS runs for 271 drugs, 60 human gut communities plus non-bacterial controls over 9 timepoints in quadruplicate to assess spontaneous drug degradation

**Table S27**: LC-MS runs for 271 drugs, 29 animal gut communities plus non-bacterial controls over 9 timepoints in quadruplicate to assess spontaneous drug degradation

**Table S32**: Reference genomes used for host DNA subtraction

**Table S33**: Overview of QC and host filtering performed with KneadData

**Table S34**: Taxonomic composition of 60 human metagenomes

**Table S35**: Taxonomic composition of 29 animal metagenomes

**Table S11**: Reference protein sequence of 31 genes known to be responsible for drug biotransformation.

**Table S13**: Quantification of reference protein marker abundance in 60 human metagenomes **Table S36**: Quantification of reference protein marker abundance in 29 animal metagenomes **Table S37**: Taxonomic composition of deposited human western metagenomes

**Table S38**: Metadata associated to the taxonomic composition of deposited human western metagenomes

**Table S39**: Samples with intensity below the noise threshold

**Table S40**: Samples with intensity below 1% of the drug average intensity in the no-bacterial control samples

**Table S41**: Outlier samples excluded from analysis

**Table S42**: Noisy drug/community combination excluded from analysis

**Table S43:** Overview of Fold Change and significance value for each community-drug-timepoint combination.

**Table S44:** List of significantly quickly biotransformed community-drug combination.

**Table S45:** List of significantly biotransformed community-drug combination.

---

**Figure S1.**
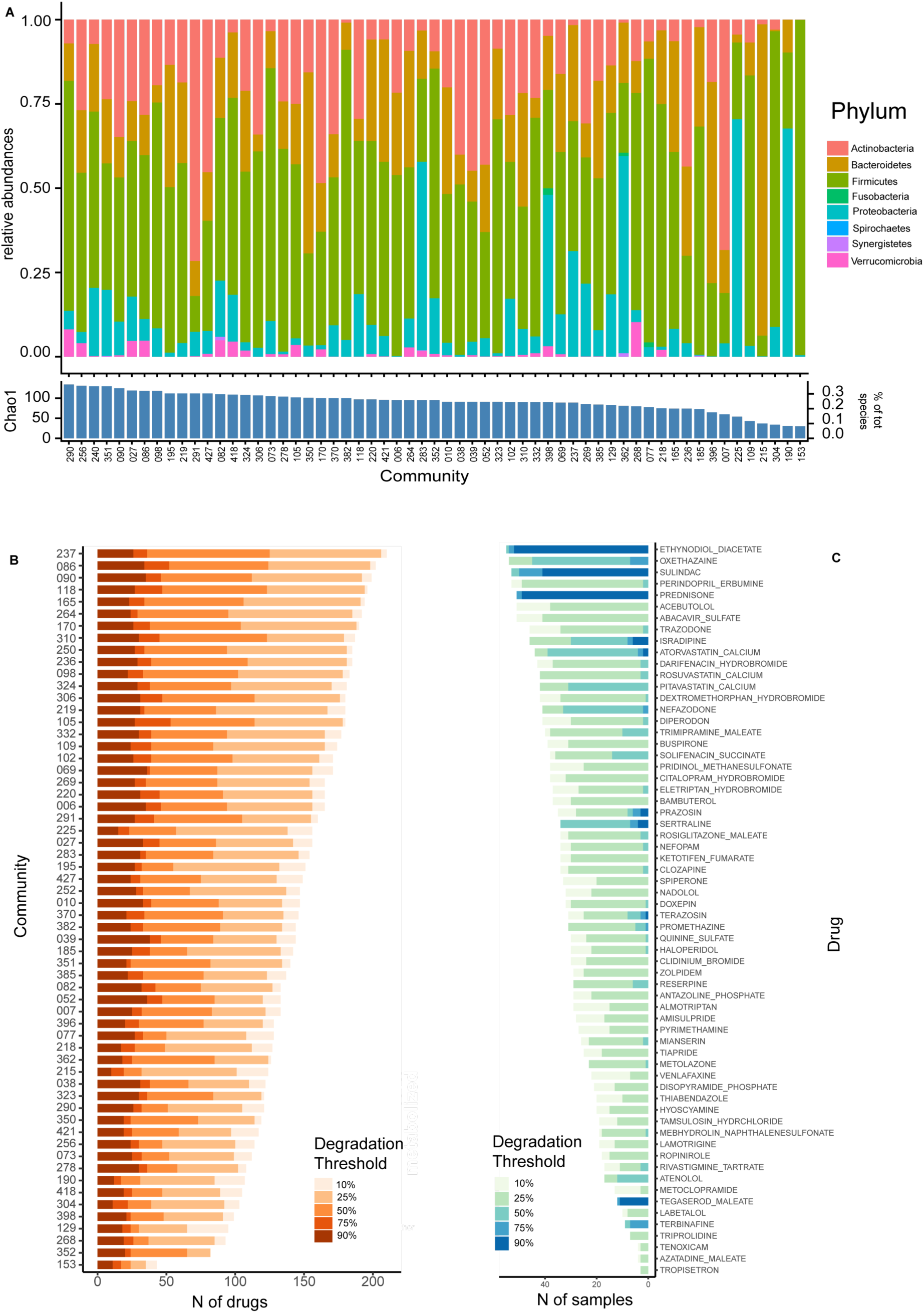
Compositional and biotransformation diversity of the tested communities and drugs. (A) Taxonomic composition (phylum level) of the 60 human-derived gut microbial communities. Lower panel indicates with a bar, for each sample, its alpha diversity value in terms of Chao1 index (left) and in terms of percentage of total species found in the whole screen (right). (B) Number of drugs metabolized by each human-derived gut microbial community, at different biotransformation thresholds. (C) For the 64 drugs that were metabolized by at least one of the 60 communities measured in this study and none of the individual species studied in Zimmermann et al.^4^, the number of communities with drug-metabolizing activity at different biotransformation thresholds is shown.

**Figure S2.**
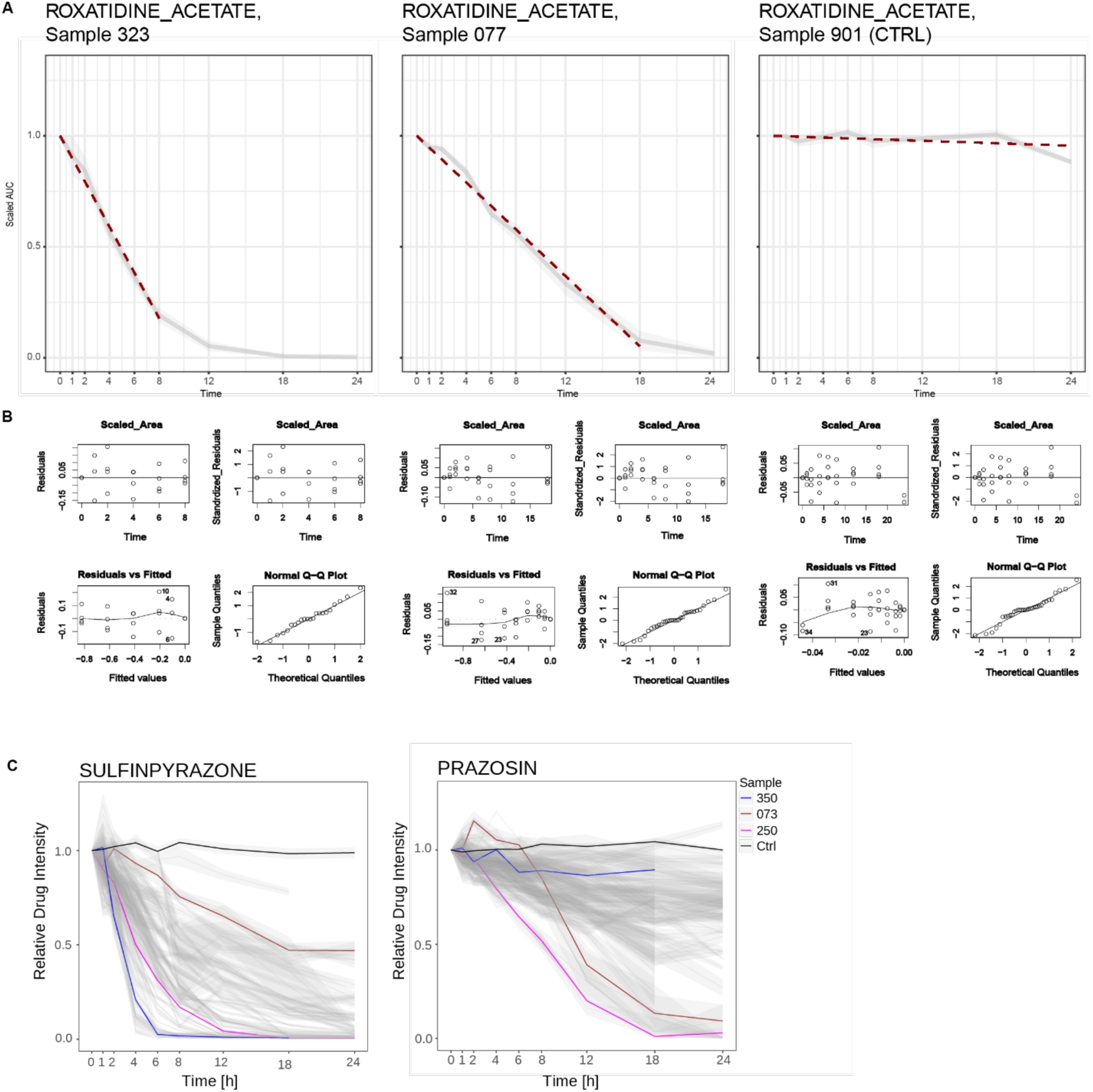
Different communities show different biotransformation rates, modelled as local linear slopes. (A) Example of the linear approximation method used for measuring drug biotransformation rates in communities, shown for the drug roxatidine acetate, two bacterial communities and a no-microbe control (Sample 901). (B) Linear modelling diagnostics for the examples in (A). (C) Drug biotransformation rates are not an inherent property of a given bacterial community. Three communities that highlight drug-specific biotransformation rates are indicated in red, blue, and magenta.

**Figure S3.**
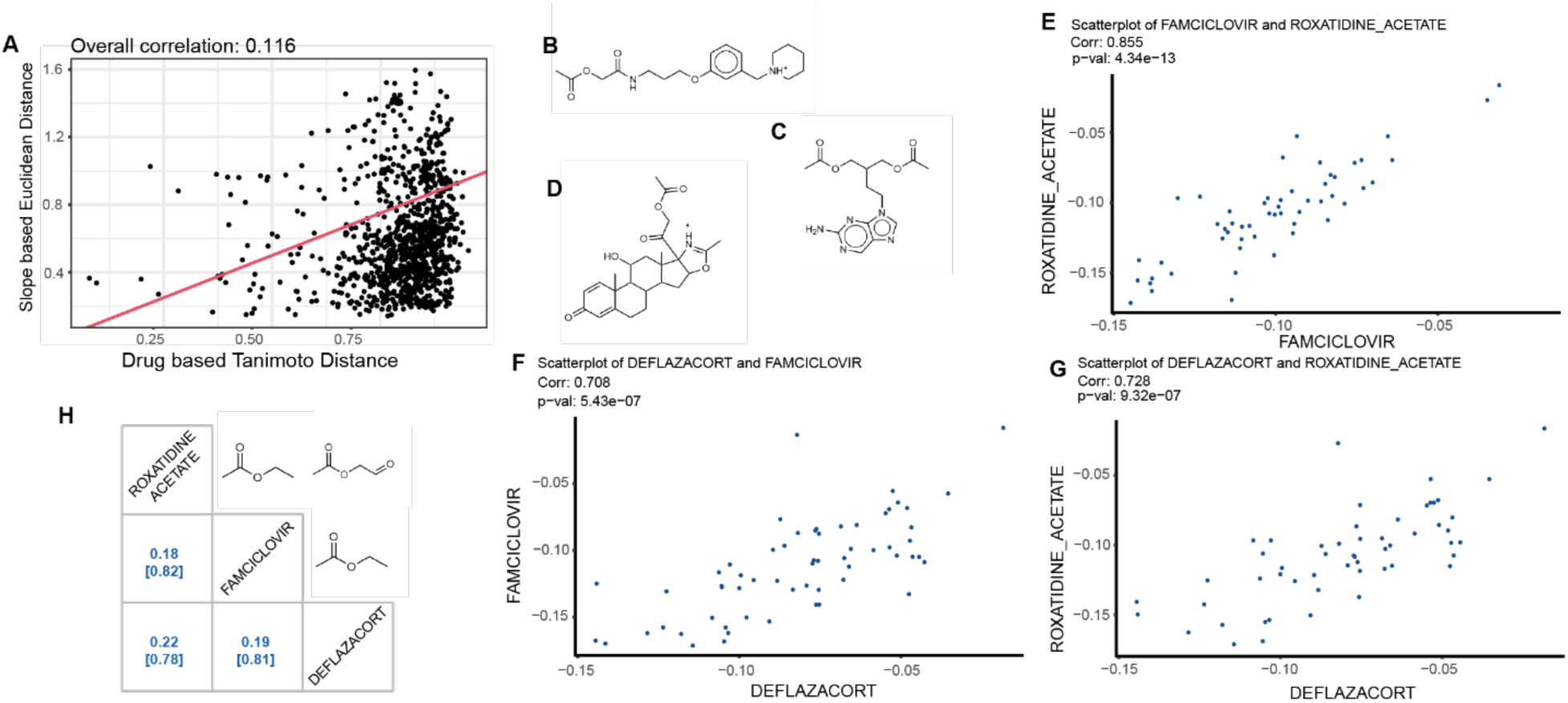
Biotransformation rates of drugs and their chemical similarity. (A) Scatter plot of chemical dissimilarity of drugs (measured as Tanimoto distance) and biotransformation rate dissimilarity (measured as Euclidean distance). The two measures do not correlate, indicating that aside from a few specific cases, overall drug structural similarity is not sufficient to determine microbiota drug transformation. (B-D) Chemical structures of roxatidine acetate (B), famciclovir (C), and deflazacort (D). (E-G) Correlation plots of biotransformation rates of famciclovir, roxatidine acetate, and deflazacort across the 60 microbial communities. (H) Mutual chemical similarity (dissimilarity below in squared brackets) of the three drugs in Fig. 2D and Fig. S3B-D, highlighting the overall low chemical similarity between these drugs despite the high and significant correlations between their biotransformation rates across communities. The upper diagonal matrix showcases the MCS (Maximum Common Substructure) between each pair of drugs.

**Figure S4.**
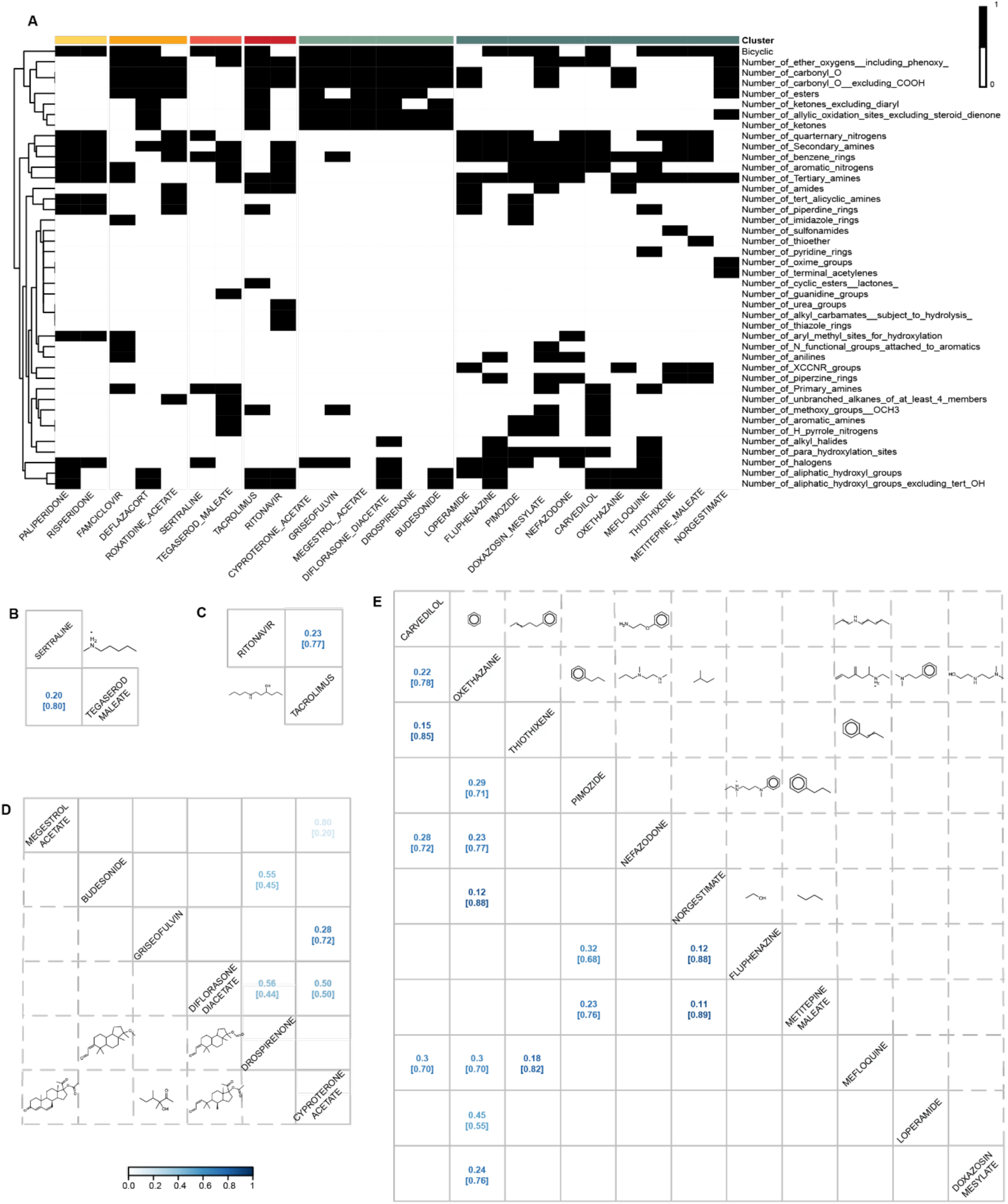
Drug biotransformation rates and their chemical similarity (continued from Figure S3). (A) Heatmap showing the presence (black) or absence (white) of specific chemical features in the top 25% entropy drugs. Drugs are clustered into groups corresponding to Figure 2B. (B-E) Matrix showing the mutual chemical similarity (dissimilarity below in squared brackets) between the drugs in panel A.

**Figure S5.**
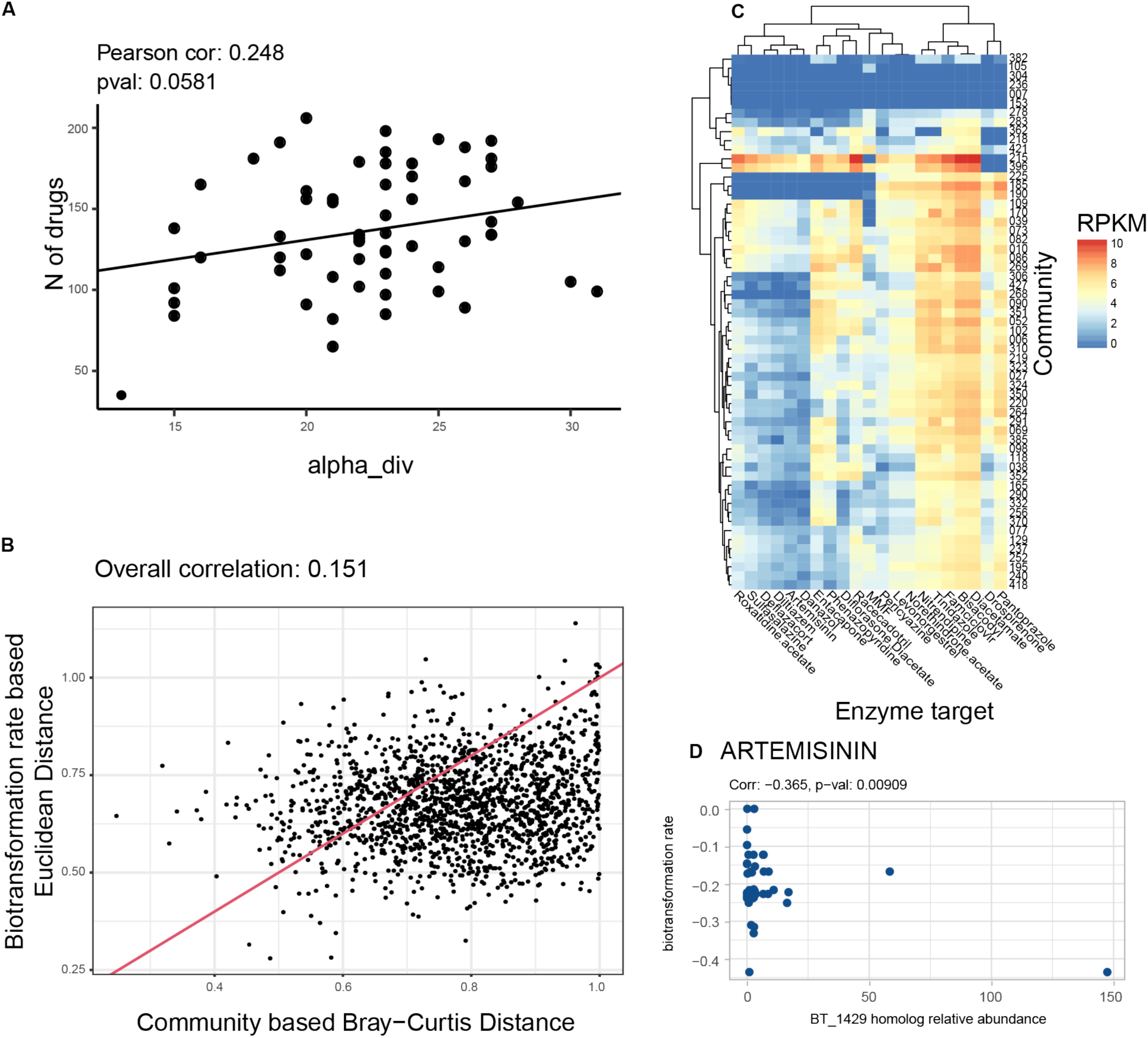
Relationships between community composition and drug biotransformation rates. (A) Scatterplot of microbiome alpha diversity (measured as Chao1) and the number of drugs biotransformed by at least 25% by the same community. (B) Scatterplot of microbiome beta diversity (measured as Bray-Curtis distance) and their dissimilarity in biotransformation rate (measured as Euclidean distance). (C) Quantification of the representation of 31 genes encoding enzymes experimentally demonstrated to be capable of metabolizing 21 of the tested drugs^4^. RPKM = reads per kilobase of reference sequence per million sample reads. (D) Scatterplot of abundance of homologs of the gene encoding BT_1429 (responsible for artemisinin metabolism) across the 60 communities and the corresponding artemisinin biotransformation rate for each community.

**Figure S6.**
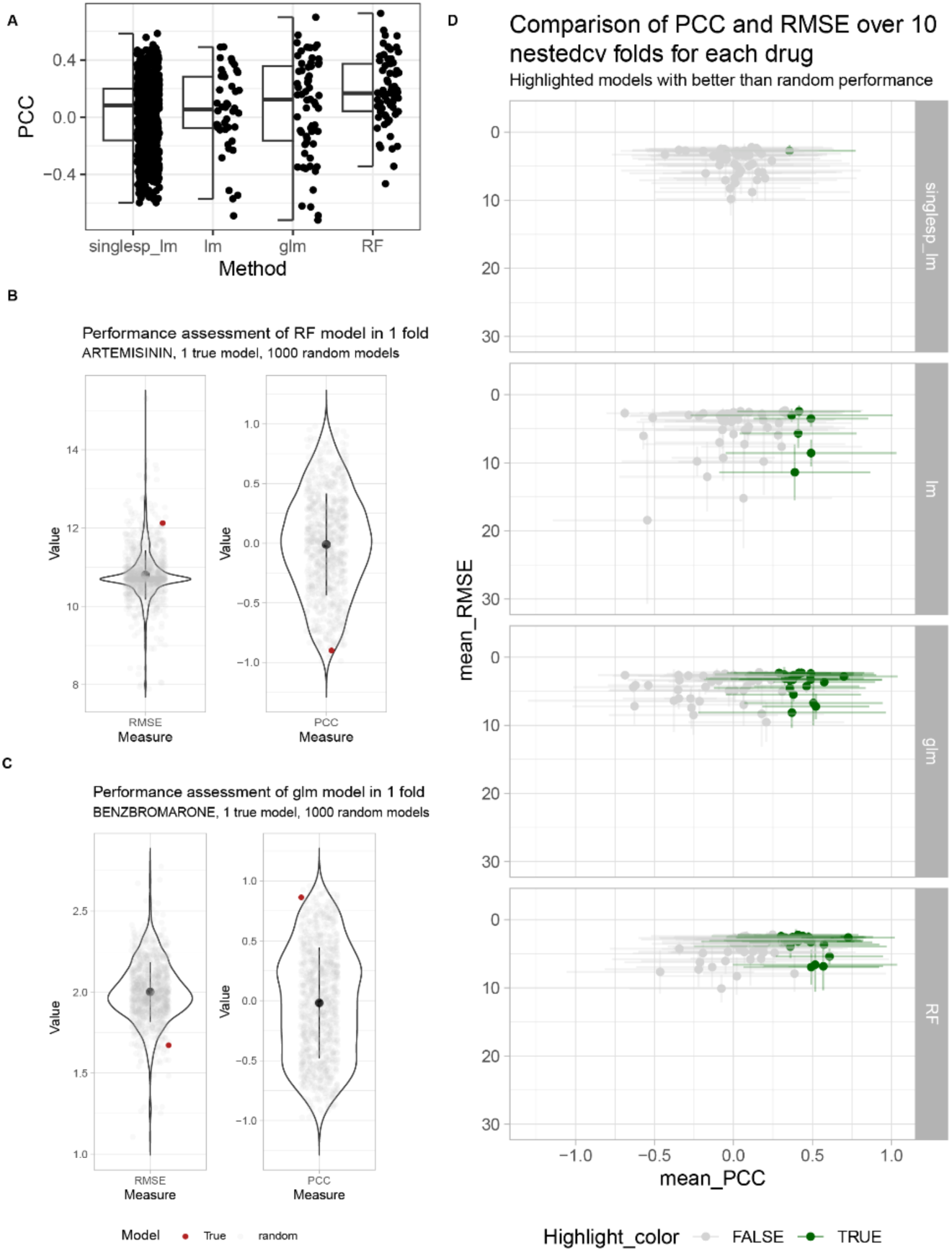
Assessment of models to predict drug biotransformation rates. (A) Average PCC (measured with nested cross-validation) between predicted and measured biotransformation rate for each of the drugs using 4 different modelling approaches: linear model using the relative abundance of individual species (singlesp_lm), combined linear models (lm), generalized linear models (glm) and random forests (RF). (B-C) Scatterplot showing the performance of the model prediction for the drug artemisinin/benzbromarone (red dot) compared to the prediction obtained by 1000 models trained on random reshuffling of the same data. This assessment was performed for 10 nested cross validation folds for PCC and RMSE, and the average performance was measured. Panel B represents a case in which the true model performs worse than random, while panel C represents a case in which the true model performs better than random. (D) Scatterplot of the mean and standard deviation of PCC and RMSE for each drug over the 10 nested cross validation folds. The models showing better than random performance are highlighted in color.

**Figure S7.**
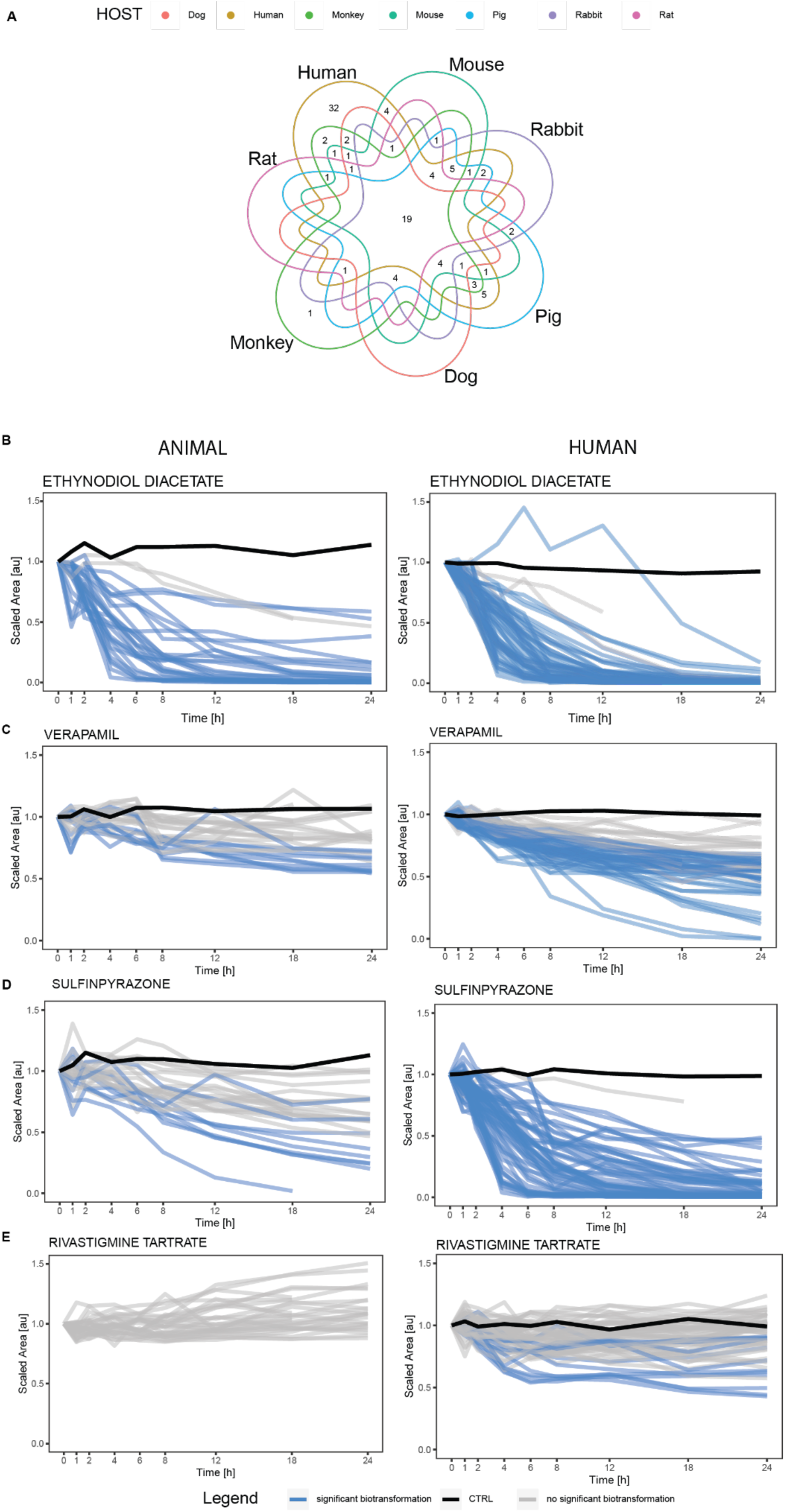
Overview of metabolomics analysis in preclinical animal models compared to humans. (B) Nested Venn diagram of efficiently metabolized drugs (greater than 75% reduction in parent drug levels) across human and animal cohorts. Only 19 of the 99 drugs that are efficiently metabolized in humans are also efficiently metabolized in all of the microbiomes from the preclinical animal models. (B-E) Case examples of drugs with similar or different biotransformation rates between human and animal samples. For some drugs (*e.g.,* ethynodiol acetate; panel B), the animal model biotransformation data recapitulates the range of biotransformation rates observed in the human cohort. In other examples, animal model biotransformation rates either only partially represent human variability in biotransformation rates (*i.e.,* verapamil; panel C), underestimate biotransformation rates (*i.e.,* sulfinpyrazone; panel D), or fail to capture metabolism altogether (*i.e.,* rivastigmine tartrate; panel E).

